# A dish-to-biobank framework links β-cell nutrient-stress programs to genetic and dietary risk for Type 2 Diabetes

**DOI:** 10.64898/2026.06.12.731989

**Authors:** Xianming Wang, Hanbyul Lee, Ann Le, Berk Turhan, Nan Hu, Pretty S. Garcia, Xuewei Cao, Dingyu Liu, Thahmina A. Ali, Nan Zhang, Breanna Williams, Caleb A. Lareau, Gao Wang, Danwei Huangfu, Kushal K. Dey

**Author notes:** Authors contributed equally (Xianming Wang, Hanbyul Lee, Ann Le, Berk Turhan).

## Abstract

Type 2 diabetes (T2D) arises from genetic susceptibility and chronic metabolic stress, but whether these converge on shared molecular programs in human populations remains unclear. Here, we develop a dish-to-biobank framework linking controlled β-cell perturbation to population-scale disease genetics through the circulating plasma proteome, and apply it to T2D. scRNA-seq of human stem cell–derived islets under factorial glucose and palmitate exposure identifies their combination (glucolipotoxicity) as the condition eliciting the strongest SC-β cell transcriptional response, with glucolipotoxicity-upregulated genes uniquely enriched for T2D heritability, monogenic diabetes genes, and rare-variant burden signals. CRISPR knockout of β-cell identity regulators *PAX6* and *PDX1* aligns with this program, establishing convergence of environmental and genetic perturbations on a shared disease-relevant state. We then used the plasma proteome as an accessible population-scale readout of these experimentally defined β-cell stress programs, scoring 45,956 UK Biobank White British participants. We define heritable stress signatures that associate with refined carbohydrate and saturated fat intake, and undergo trans-tissue genetic regulation, with a subset of variants showing diet-dependent effects. Together, these findings establish glucolipotoxicity as a genetically anchored model of β-cell dysfunction and provide a generalizable framework for linking controlled cellular perturbations to human disease genetics at population scale.

## Introduction

Complex diseases are difficult to mechanistically resolve because inherited susceptibility and environmental exposure are distributed across tissues, time, and physiological state, while the relevant human cell states are often inaccessible in living individuals. Genome-wide association studies have identified thousands of loci for complex diseases and traits^1–3^, and single-cell genomic approaches have begun to link these variants to specific cell types, cellular states, and regulatory gene programs^4–8^. Yet how these cellular programs connect to disease genetic architecture and environmental exposures which raise disease risk remains less well understood. Type 2 diabetes (T2D) offers a compelling case to address these gaps: it arises from the interplay between genetic susceptibility and chronic metabolic stress, manifesting through systemic insulin resistance and progressive pancreatic β-cell dysfunction^9–11^. Although obesity, chronic hyperglycemia, dyslipidemia, and inflammation are well-established drivers of disease progression, the specific cellular programs through which diabetogenic environments impair β-cell function remain incompletely defined^12^. Nutrient excess — particularly the chronic combination of elevated glucose and saturated fatty acids such as palmitate, commonly referred to as glucolipotoxicity — has long been implicated in β-cell failure^13,14^; however, whether it represents a distinct molecular insult — or simply the additive combination of glucotoxic and lipotoxic effects — and how the resulting programs relate to human disease risk, remains an open question.

While early bulk transcriptomic studies provided initial insights into islet responses to glucose and fatty acids^15,16^, single-cell RNA-seq (scRNA-seq) has since enabled cell-type-specific dissection of stress responses in human islets — including profiling of high-glucose exposure^17^ and a systematic screen of endoplasmic reticulum (ER) and inflammatory stressors^18^. However, no study has systematically compared glucotoxic, lipotoxic, and combined glucolipotoxic stress at single-cell resolution in human β cells using a factorial design. Critically, whether nutrient stress–induced transcriptional programs correspond to the genes and pathways implicated by human T2D genetics remains unknown. Population cohorts capture real-world genetic, dietary, molecular, and disease variation at scale, but are observational, noisy, and weak on mechanism; controlled cellular systems provide causal precision and cell-type resolution, but simplify the chronic, multi-organ physiology of disease. Bridging this gap requires a framework that connects causal perturbations in controlled in vitro systems with the genetically and environmentally complex landscape of human populations, where variation in genotype and dietary exposure can be measured at scale.

We reason that stress-induced gene programs in pancreatic endocrine cells — which release their products, including insulin, glucagon, and stress-responsive proteins, directly into the bloodstream — should leave detectable molecular signatures in the circulating plasma proteome. This motivates a “dish-to-biobank” framework in which the plasma proteome serves as a bridge spanning three interconnected layers: (i) β-cell transcriptional programs induced by nutrient and genetic perturbations in controlled in vitro systems; (ii) projection of these programs onto the plasma proteome as secreted stress signatures measurable in human populations; and (iii) multi-tissue genetic and environmental regulation of these signatures, including contributions from hepatic and adipose regulators of the circulating nutrient milieu and dietary intake (**Figure 1**). The framework is now testable through the convergence of two complementary resources: large-scale plasma proteomic, metabolomic and genetic data from a population cohort^19,20^, and human pluripotent stem cell (hPSC)–derived islet models^21–23^ that provide a controlled in vitro platform for generating the causal molecular programs.

**Figure 1:**
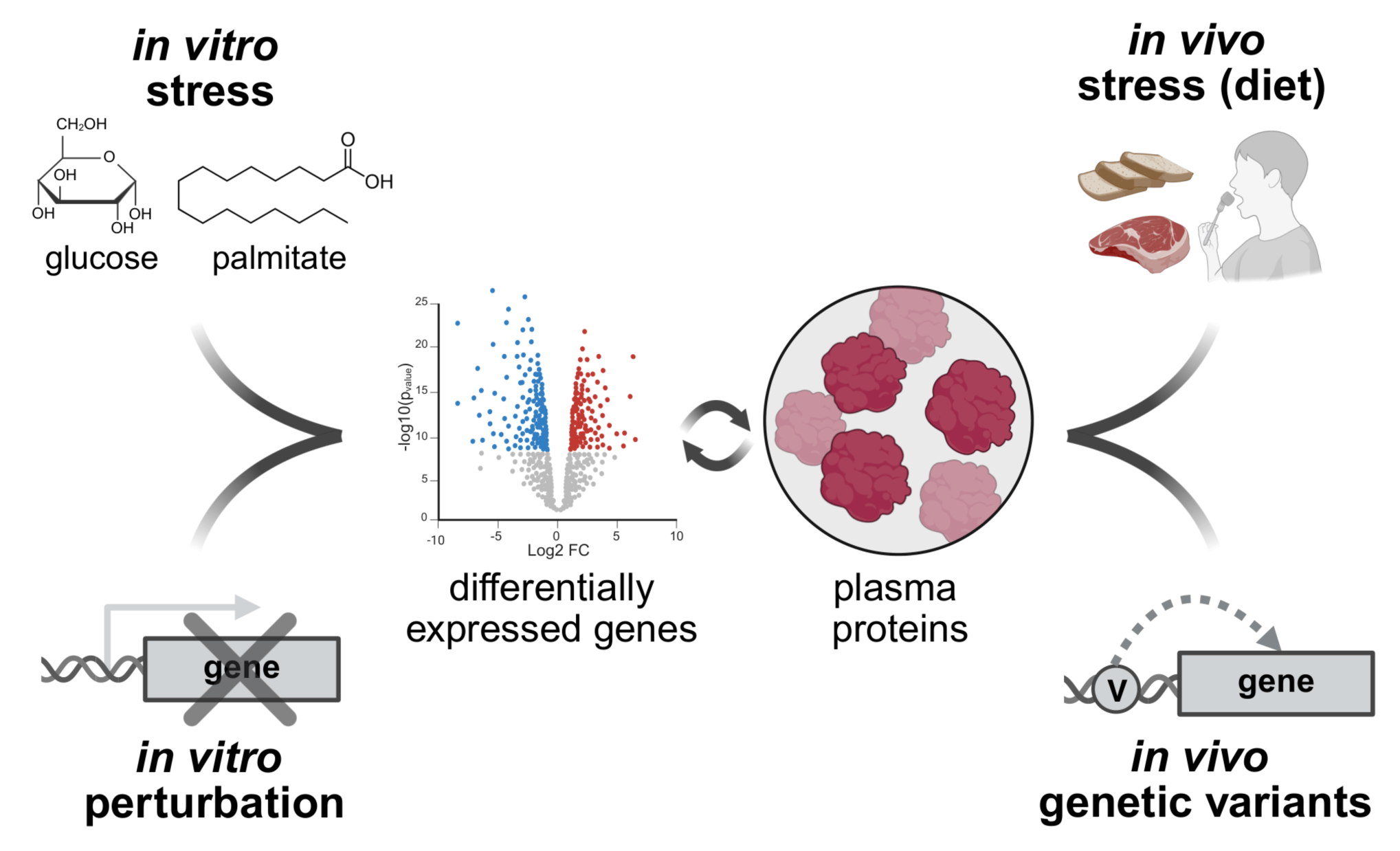
Overview of the dish-to-biobank framework. Stress-responsive gene programs defined in stem cell–derived islets under glucolipotoxic stress (top left) and supported by CRISPR perturbation (bottom left) are projected onto population-scale plasma protein abundance (center), which serves as a bridge linking *in vitro* mechanism to *in vivo* dietary exposure (top right) and common genetic variation (bottom right) that shape human disease risk. Thus, the framework links causal but simplified cellular perturbations to noisy but physiologically and genetically rich human population data.

Here, we systematically model nutrient stress in human stem cell–derived islet clusters using a factorial design of glucose and palmitate exposure, coupled with scRNA-seq. We show that glucolipotoxicity elicits the most pronounced and disease-relevant transcriptional response in SC-β cells, characterized by increased insulin gene expression alongside activation of ER stress pathways. By integrating these in vitro causal programs with common and rare variant genetic data, in vivo diabetes mouse models^24^, transcriptional changes in human T2D islets^25,26^, and CRISPR loss-of-function perturbation datasets^27^, we demonstrate that glucolipotoxicity captures a uniquely informative dimension of β-cell dysfunction that converges across the allele-frequency and disease-severity spectrum, from common population risk for T2D to monogenic forms of diabetes. To connect these cellular programs with human physiology, we project stress-induced gene sets onto the plasma proteome of 45,956 UK Biobank White British participants^20^, revealing circulating stress signatures that are heritable, associate with dietary patterns, and colocalize with T2D-associated loci. Together, our study establishes a generalizable framework linking controlled in vitro perturbations to population-scale disease genetics through circulating biomarkers. Our findings identify glucolipotoxicity as a genetically anchored model of β-cell dysfunction and suggest that environmental nutrient stress and genetic perturbation of β-cell identity regulators converge on shared molecular programs underlying T2D pathogenesis.

## Results

### Single-cell profiling of SC-islets reveals coordinated islet response to glucolipotoxicity

Stem cell–derived islet-like clusters provide a powerful platform to interrogate how environmental and genetic perturbations shape human β-cell biology. To leverage this system, XM001 induced pluripotent stem cells (iPSCs)^28^ were differentiated into islet-like clusters and subjected to a 2×2 factorial experimental design combining glucose and palmitate exposure to model key diabetogenic nutrient conditions (**Figure 2a**). Specifically, islet clusters were exposed for 72 hours to a factorial combination of glucose (8 mM basal vs 33 mM high) and palmitate (1 mM palmitate–BSA vs BSA vehicle control), enabling comparison of glucotoxic, lipotoxic, and combined glucolipotoxic conditions (**Methods**). SC-islets were profiled using high-throughput scRNA-seq across all conditions, yielding 38,350 high-quality cells (382,245,713 UMIs) after quality control **(Methods)**. Unsupervised Louvain clustering identified 8 transcriptionally distinct cell populations annotated using canonical cell-identity markers for SC-α, SC-β, SC-δ, SC-enterochromaffin, polyhormonal, proliferating α, mesenchyme, and ductal progenitor cells (**Figure 2b,c**), rather than stress conditions (**Supplementary Figure 1a, Supplementary Table S1)**.

**Figure 2:**
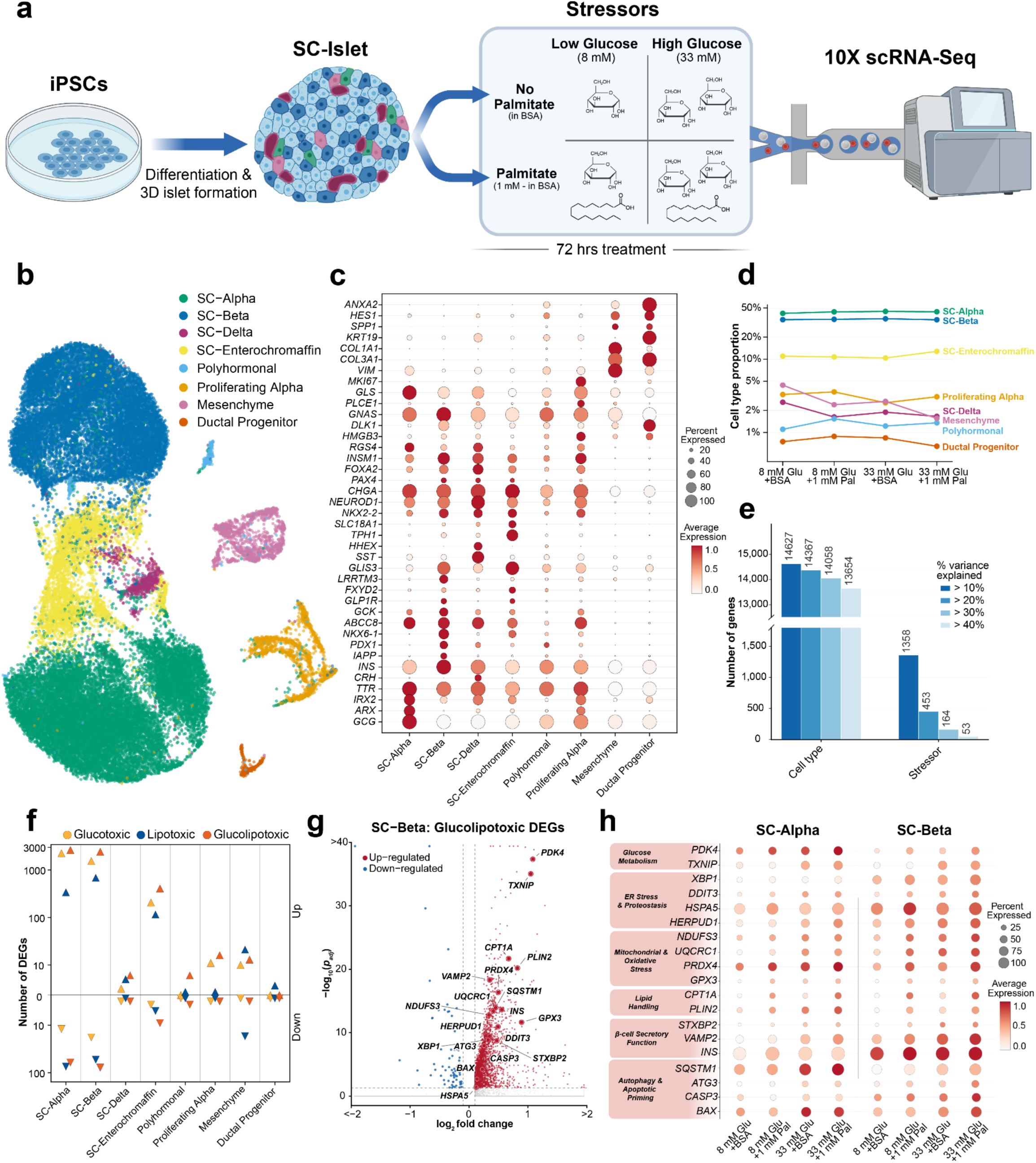
Single-cell RNA sequencing of nutrient-stressed SC-islets. **a.** Schematic of the experimental design: iPSC-derived islets were exposed for 72 h to a 2×2 factorial combination of glucose (8 mM baseline, 33 mM high) and palmitate (1 mM palmitate–BSA vs BSA vehicle) and profiled by 10X single-cell RNA-sequencing (N = 38,350 cells). **b.** Uniform manifold approximation and projection (UMAP) of islet cells across all treatment conditions, colored by the eight annotated cell types. **c.** Dot plot of canonical marker-gene expression across annotated cell types. Dot size indicates the percentage of cells expressing the gene; color indicates average expression magnitude. **d.** Proportions of all annotated SC-islet cell types across the four nutrient-stress conditions. **e.** Variance component analysis across all genes. Bars show the number of genes for which cell type versus stress condition explains >10%, >20%, >30%, or >40% of expression variance; cell type accounts for the largest fraction, followed by stress condition. **f.** Number of up- and down-regulated DEGs per cell type under glucotoxic (33 mM Glu + BSA vs 8 mM Glu + BSA), lipotoxic (8 mM Glu + 1 mM Pal vs 8 mM Glu + BSA), and glucolipotoxic (33 mM Glu + 1 mM Pal vs 8 mM Glu + BSA) conditions. DEGs were defined as adjusted P < 0.05 and |log₂FC| > 0.1 (test and correction as in g). **g.** Volcano plot of differential expression in SC-β cells under glucolipotoxicity relative to the baseline condition (8 mM glucose + BSA vehicle). Differential expression was assessed by Wilcoxon rank sum test with Bonferroni correction across all genes as implemented in Seurat; dashed lines mark the significance thresholds (adjusted P < 0.05, |log₂FC| > 0.1). **h.** Dot plot of selected genes in SC-α and SC-β cells across the four nutrient-stress conditions, grouped by functional category (glucose metabolism, ER stress and proteostasis, mitochondrial and oxidative stress, lipid handling, β-cell secretory function, and autophagy/apoptosis-related genes). Dot size indicates the percentage of cells expressing the gene; color indicates average expression magnitude. Numerical results are reported in **Supplementary Table S1**.

Across conditions, we observed consistent proportions of each cell type, indicating that nutrient stress has minimal impact on islet cell composition (**Figure 2d**). Expectedly, cell cycle differences accounted for clear sub-populations among proliferating cell types, but not in other cell types (**Supplementary Figure 1b**). Variance component analysis across all genes confirmed that cell type explains the largest fraction of transcriptional variation, with stress condition emerging as the second-largest source — affecting >10% of variance for 1,358 genes (**Figure 2e**) — motivating cell type–specific analyses of stress response. Pairwise differential expression analysis relative to baseline (8 mM glucose with BSA vehicle; Wilcoxon rank sum test, adjusted P < 0.05, |log₂FC| > 0.1) revealed that the combination of high glucose and palmitate (glucolipotoxicity) induced the largest transcriptional response in SC-β, SC-α, and SC-enterochromaffin cells, with substantially more up-regulated genes (SC-β: 2,423; SC-α: 2,629; SC-enterochromaffin: 407) than down-regulated genes (SC-β: 78; SC-α: 61; SC-enterochromaffin: 9) (**Figure 2f,g, Supplementary Table S1**). This directional asymmetry is consistent with an active transcriptional response rather than transcriptional collapse. We next focused on the two major endocrine cell types, SC-α and SC-β, to examine shared and cell type–specific responses to glucolipotoxic stress. Among the genes up-regulated in both SC-α and SC-β cells under glucolipotoxicity were *PDK4* and *TXNIP*, well-established regulators of glucose metabolism and stress responses^29–31^ **(Figure 2g,h)**. We identified 1,308 genes jointly up-regulated in both cell types, enriched for pathways related to mitochondrial respiration and oxidative phosphorylation. Cell type–specific programs were dominated by cytoplasmic translation and ribosomal function in SC-α (1,321 genes) versus protein processing and N-linked glycosylation in SC-β (1,115 genes) (**Supplementary Figure 1c**).

Focusing on SC-β cells, glucolipotoxicity induced a coherent transcriptional program spanning ER stress/proteostasis (i.e., maintenance of protein folding and quality control; e.g., *XBP1*, *DDIT3*, *HSPA5*, *HERPUD1*), mitochondrial/oxidative stress (e.g., *NDUFS3*, *UQCRC1*, *PRDX4*, *GPX3*), and lipid handling (*CPT1A*, *PLIN2*), reflecting coordinated metabolic remodeling (**Figure 2h**). Activation of vesicle trafficking machinery (*STXBP2, VAMP2*) and increased *INS* expression were also observed (**Figure 2h**). These changes are consistent with a compensatory β-cell secretory program. Downstream of these changes, stress-response pathways including autophagy (*SQSTM1*, *ATG3*) and apoptotic regulators (*CASP3*, *BAX*) were induced, without evidence of executioner apoptosis or loss of β-cell abundance. Together, these features suggest an early stress-responsive state characterized by increased expression of genes involved in β-cell secretory function, which may reflect elements of a compensatory response.

### Nutrient-induced stress programs are uniquely informative for T2D heritability

To identify which cellular stress programs are most relevant to the genetic architecture of T2D, we quantified the contribution of stress-induced gene sets to common-disease heritability. We related GWAS signals for diabetes-related traits (average GWAS sample size = 245,580; **Supplementary Table S2)**^32–34^ to differentially expressed gene (DEG) sets from our 3 nutrient-stress states (glucotoxicity, lipotoxicity, glucolipotoxicity). Using sc-linker^4^, we constructed variant-level annotations by mapping genes in each program to regulatory elements via the ENCODE-rE2G method^35^, and evaluated disease relevance using two complementary metrics: heritability enrichment (ENR) and standardized effect size (*τ*\*) conditional on 97 baseline-LD (v2.2) coding, conserved, and LD-related annotations^36,37^ (**Methods**). When meta-analyzed across nutrient stressors, diabetes-related traits (T2D, T1D, HbA1c, fasting glucose) showed the strongest enrichment in SC-α and SC-β cells (T2D: ENR = 2.5 and 2.9, respectively) (**Supplementary Figure 2a**); we therefore focused downstream analyses on SC-α and SC-β cells.

Within this endocrine compartment, we next asked which specific stress program is most T2D-informative, evaluating the DEG sets from our 3 nutrient-stress conditions alongside seven additional primary β-cell and α-cell stress states — 2 ER-stress and 5 inflammatory — from an external single-cell screen^18^, which together define a complementary stressor taxonomy. In β cells, glucolipotoxicity and glucotoxicity up-regulated DEGs (uDEGs) showed the strongest heritability enrichments for T2D (ENR = 4.2, p= 3e-03 and 4.4, p= 4e-03, respectively), whereas lipotoxicity uDEGs did not reach significance (**Figure 3a**). Notably, glucolipotoxicity and glucotoxicity uDEGs were the only gene programs showing nominally significant (p < 0.05) *τ*\* for T2D (**Figure 3b**); in a joint model, glucolipotoxicity absorbed the signal from glucotoxicity and remained the only jointly significant program (**Supplementary Figure 2b**). In α cells, nutrient-stress uDEGs and dDEGs did not reach significance for T2D *τ*\* (**Supplementary Figure 2c**), indicating that β-cell dysfunction contributes more substantially to T2D heritability than α-cell dysfunction under nutrient stress and motivating our focus on β-cell programs in subsequent analyses. We next asked whether disease relevance differed across stressor classes. Stressors showed divergent T2D and T1D associations consistent with known disease etiology: nutrient- and ER-stress uDEGs exhibited 2.6-fold higher T2D enrichment on average compared to inflammatory-stress uDEGs, whereas inflammatory-stress uDEGs showed 1.9-fold higher T1D enrichment (**Figure 3a**). Neither ER-stress nor inflammatory-stress uDEGs showed significant *τ*\* for T2D or T1D, though inflammatory-stress uDEGs showed a suggestive association with T1D (p < 0.15) (**Figure 3b**). Consistent with this divergent disease enrichment, unsupervised clustering of stressors by effect-direction concordance (EDC)^38^grouped the nutrient and canonical ER stressors (brefeldin A, thapsigargin) together and separated them from the inflammatory perturbations (within-cluster EDC = 88% for the nutrient/ER cluster; 98% for the inflammatory cluster; cross-cluster EDC = 60%) (**Figure 3c**). Taken together, T2D risk concentrates in metabolic/nutrient stress pathways, T1D risk in inflammatory pathways, and glucolipotoxicity occupies the most T2D-informative position within the metabolic axis. Thus, human genetic architecture allows prioritization of specific stress responses most likely to lie on causal disease-relevant pathways in people.

**Figure 3:**
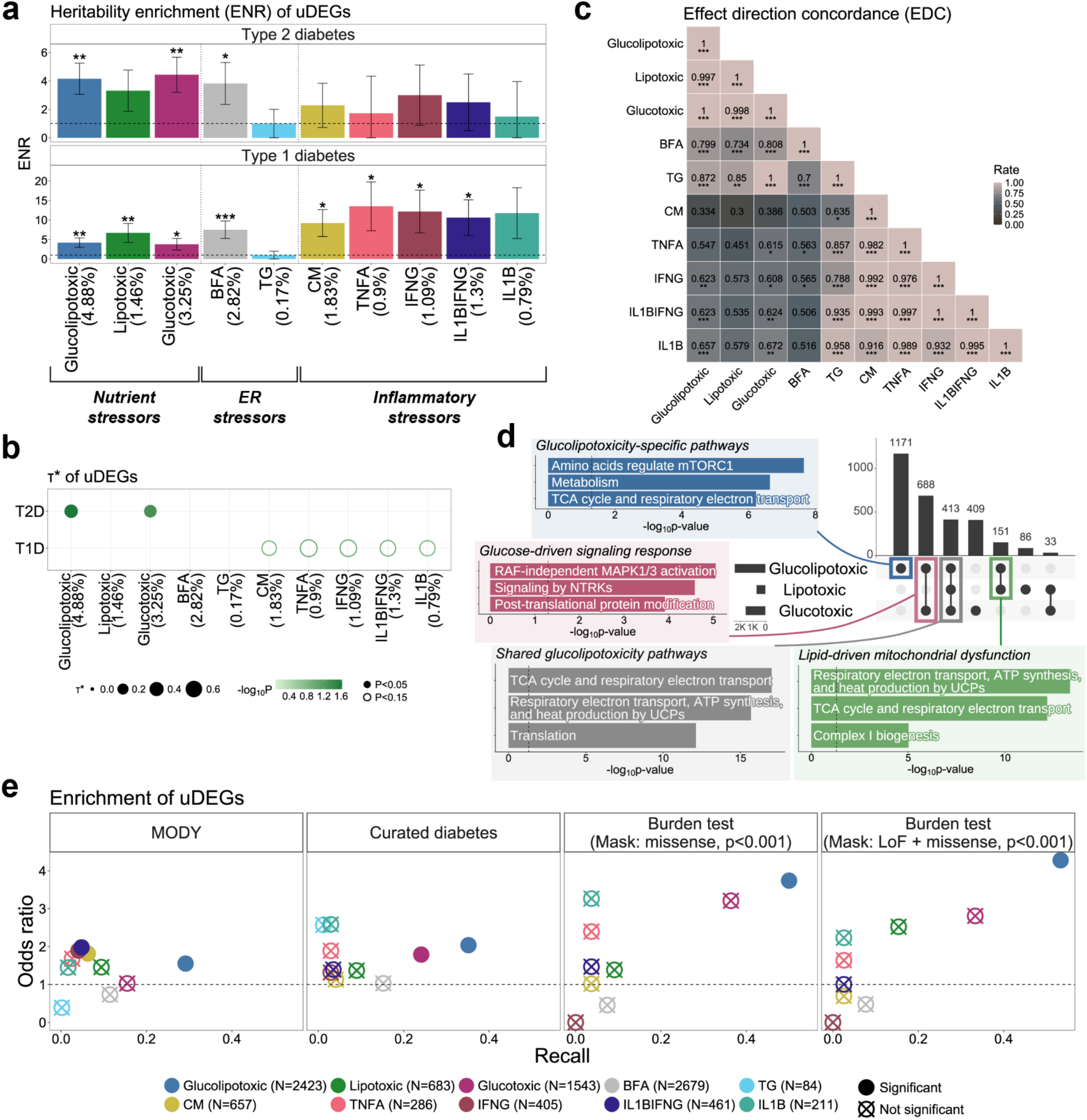
Comparative analysis of disease-relevant genetic signals across stress-induced signatures in SC-β cells. **a.** Heritability enrichment (ENR) of up-regulated differentially expressed genes (uDEGs) under three nutrient stressors (glucolipotoxicity, lipotoxicity, and glucotoxicity), two external ER stressors (BFA: brefeldin A; TG: thapsigargin), and five external inflammatory stressors (CM: IL1β + IFNγ + TNFα; TNFA: TNFα; IFNG: IFNγ; IL1BIFNG: IL1β + IFNγ; IL1B: IL1β) for Type 2 Diabetes (T2D) and Type 1 Diabetes (T1D). Error bars represent ±1 SE. The dashed line marks ENR = 1. The percentage in parentheses below each stressor name indicates the proportion of common variants falling within the regulatory annotation (linked enhancers) of its uDEG set. **b.** Standardized effect sizes (*τ*\*) of uDEGs under the same 10 stressors for T2D and T1D. Dot size represents the magnitude of *τ*\*, dot color represents significance (−log_10_P), filled dots denote P < 0.05 and open dots P < 0.15. **c.** Effect direction concordance (EDC) of DEGs between each pair of the 10 stressors (**Methods**). Numbers and shading within each cell indicate the EDC rate. **d.** UpSet plot of uDEGs under glucolipotoxicity, lipotoxicity, and glucotoxicity, with the top enriched pathways for the 1,171 glucolipotoxicity-specific uDEGs (blue), the 688 uDEGs shared exclusively between glucolipotoxicity and glucotoxicity (pink), the 151 uDEGs shared exclusively between glucolipotoxicity and lipotoxicity (green), and the 413 uDEGs shared across all three nutrient stressors (grey). **e.** Enrichment of stressor uDEGs against three classes of diabetes validation gene sets - (i) 561 MODY-associated genes from ref^44^, (ii) 175 expertly curated genes associated with monogenic and polygenic forms of diabetes (**Methods**), and (iii) genes reaching FDR < 0.1 in rare-variant burden tests for HbA1c across two coding masks (missense and LoF + missense; MAF < 0.001). Recall (x-axis) is the fraction of validation-set genes recovered as uDEGs under each stressor; the odds ratio (y-axis) and its significance are from a one-sided Fisher’s exact test on the 2×2 table cross-classifying genes by validation-set membership and uDEG status. Filled dots indicate significant overlap, crossed dots non-significant; N in the legend gives the number of uDEGs per stressor. Asterisks in panels **a** and **c** denote significance from a one-sided Z-test and one-sided exact binomial test assessing whether ENR > 1 and EDC > 0.5, respectively (*** P < 0.001, ** P < 0.01, * P < 0.05). All analyses are for β cells (SC-β for our nutrient stress and primary β cells for ER and inflammatory stress). Numerical results are provided in **Supplementary Table S2**.

We next asked why glucolipotoxicity is more T2D-informative than either of its component stressors alone. uDEGs unique to glucolipotoxicity in SC-β cells were enriched for metabolic pathways including general metabolism, mTORC1 signaling, and the TCA cycle and respiratory electron transport (**Figure 3d**), consistent with the synergistic convergence of excess glucose and lipid substrates at shared metabolic nodes. uDEGs shared exclusively between glucolipotoxicity and glucotoxicity were enriched for NTRK signaling, RAF-independent MAPK1/3 activation, and post-translational protein modification pathways (**Figure 3d**), suggesting dysregulation of growth factor receptor and kinase signaling cascades under conditions of chronic glucose excess. In contrast, uDEGs shared exclusively between glucolipotoxicity and lipotoxicity were enriched for respiratory electron transport, TCA cycle disruption, and Complex I biogenesis (**Figure 3d**), consistent with the established role of saturated fatty acids in impairing β-cell mitochondrial function^13^. uDEGs shared across all three nutrient stressors were enriched for similar pathways — respiratory electron transport and TCA cycle disruption — suggesting that mitochondrial stress represents a convergent response to nutrient excess regardless of stressor identity. Glucolipotoxicity thus combines a synergistic metabolic activation arm, a glucose-driven dysregulation arm, a lipid-driven mitochondrial stress arm, and a shared mitochondrial stress arm — each potentially contributing distinct aspects of T2D pathogenesis. We assessed the T2D heritability informativeness of gene programs corresponding to these 4 arms of glucolipotoxicity, both marginally and in a joint model; despite the convergent nature of the shared arm, the glucose-driven arm exhibited the strongest heritability information (**Supplementary Figure 2d**). Within this glucose-driven arm (glucolipotoxicity–glucotoxicity shared program), the top T2D-informative genes prioritized by MAGMA^39^ were *WFS1* (rank 6) and *RNF43* (rank 25). *WFS1* encodes an ER-resident protein that regulates the unfolded protein response and calcium homeostasis, thereby protecting β-cells from ER stress^40^. *RNF43* encodes an E3 ubiquitin ligase that negatively regulates Wnt/β-catenin signaling^41^, and may influence β-cell survival under metabolic stress. Within the glucolipotoxicity–lipotoxicity shared program, the top-ranked genes were *CISD2* (rank 23) and *UQCR10* (rank 36). *CISD2* encodes a mitochondria-associated ER protein that maintains mitochondrial integrity and redox balance under metabolic stress^42^, while *UQCR10* encodes a subunit of mitochondrial respiratory chain complex III that supports oxidative phosphorylation and cellular energy metabolism^43^ (**Supplementary Table S2**).

We next assessed whether this common-variant enrichment extends across the allele-frequency spectrum by comparing stressor programs against three classes of diabetes silver-standard gene sets: (i) 561 Maturity-Onset Diabetes of the Young (MODY) associated genes^44^, (ii) 175 expertly curated genes associated with monogenic and polygenic forms of diabetes (**Supplementary Table S2**), and (iii) genes reaching FDR < 0.1 in rare-variant burden tests for glycated hemoglobin (HbA1c) across seven coding masks (MAF < 0.001; N = 2-39 genes per mask) (**Methods**). Although the glucolipotoxicity uDEG set is larger than those of the other stressors, it showed both considerably higher recall of the silver-standard gene sets and comparable-to-higher odds ratios (OR = 1.55–4.28, P < 0.05) across MODY genes, the curated diabetes set, and two of the seven burden-test masks (**Figure 3e**, **Supplementary Table S2**) — breaking the usual trade-off between gene-set size and enrichment. Notably, these silver-standard sets showed limited overlap with one another and with MAGMA-based T2D risk genes (average overlap= 9.9%, **Supplementary Figure 2e**), indicating that they represent largely independent slices of diabetes genetic architecture. Their joint convergence on the glucolipotoxicity program therefore suggests that this program captures core β-cell dysfunction mechanisms operating across allele frequencies and disease severity.

Together, these analyses — spanning common-variant heritability, rare-variant burden signals, and monogenic diabetes genes — identify glucolipotoxicity, and specifically its glucose-driven arm, as the most informative and physiologically relevant model of the β-cell stress response underlying T2D genetic architecture.

### Genetic and nutrient stress perturbations converge on β-cell dysfunction programs that better recapitulate human T2D than mouse models

To test whether genetic perturbation of β-cell regulators recapitulates the glucolipotoxic transcriptional program — and to evaluate how both perturbation classes align with transcriptional states observed in human and mouse T2D models — we compared glucolipotoxicity DEGs against CRISPR-knockout (KO) transcriptional profiles from perturbations of 30 genes in SC-islets (28 passing QC; N = 2,966 SC-β cells)^27^, targeting genes implicated in neonatal diabetes, MODY, and islet development, and further benchmarked both perturbation classes against primary β cell differential expression between T2D and healthy islets from the Human Pancreas Analysis Program^25^ (HPAP) and against the db/db mouse model of genetic T2D^24^ (**Figure 4a**; **Methods**, **Supplementary Table S3**). 9 KO perturbations showed significant concordance of DEGs with glucolipotoxicity (FDR < 5%), with *PAX6* KO exhibiting the strongest alignment (EDC = 0.89) (**Figure 4b**). Raw KO DEGs often capture broader transcriptional changes not relevant to T2D; we therefore employ the PerturbScape method (Turhan*, Lee* in preparation; Methods) to identify T2D meta-program genes underlying each KO. *PAX6* KO and *PDX1* KO T2D meta-program genes specifically showed significant overlap with glucolipotoxic uDEGs (**Figure 4c**). This indicates convergence on T2D-relevant pathways between glucolipotoxicity and *PAX6/PDX1* KO. This convergence is bidirectional: glucolipotoxicity itself downregulates both *PAX6* and *PDX1* (log₂FC = −0.11 and −0.16, respectively), consistent with prior reports of *PAX6* and *PDX1* protein-level reduction under palmitate and glucolipotoxic exposure in rodent β-cell models and primary human islets^45–47^. We compared the top 100 T2D meta-program genes of *PAX6* KO and *PDX1* KO against glucolipotoxicity uDEGs to resolve shared versus perturbation-specific components (**Figure 4d**). Glucolipotoxicity-specific uDEGs highlight coordinated metabolic adaptation, including lipid sensing (*FFAR1*), insulin signaling (*IRS2*), and oxidative stress response (*GPX3*). In contrast, *PDX1* and *PAX6* loss—independent of glucolipotoxicity—preferentially reveals transcription factor–dependent cell identity programs, encompassing cell adhesion (*ALCAM*), extracellular signaling (*CRISPLD1*), and GPCR-mediated signal transduction (*GNG2*), consistent with lineage-specific regulatory control rather than stress adaptation. Notably, uDEGs shared between glucolipotoxicity and *PDX1/PAX6* perturbations further define a convergent remodeling program, including calcium-dependent transcription (*NFATC2*) and vesicle-associated machinery (*NRXN3*), highlighting both shared and condition-specific components of the β-cell stress response.

**Figure 4:**
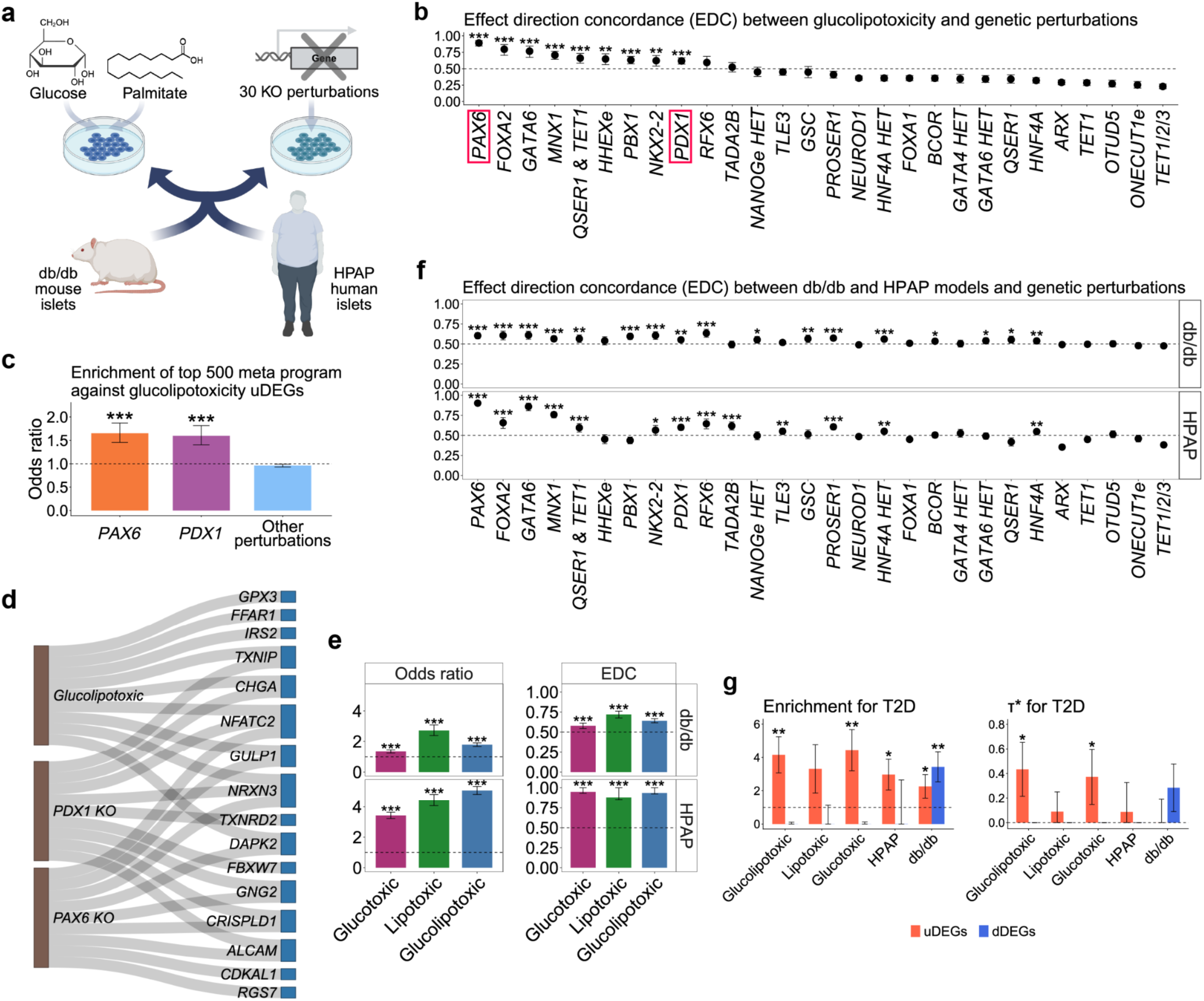
Nutrient stress and genetic perturbations converge on β-cell dysfunction programs that recapitulate human T2D transcriptional architecture. **a.** Schematic of the comparative framework. Glucolipotoxic nutrient stress (glucose + palmitate) and CRISPR-knockout (KO) perturbations of 30 genes in SC-islets were benchmarked against transcriptional profiles from human T2D islets (HPAP) and the db/db mouse model. **b.** Effect direction concordance (EDC) between glucolipotoxicity DEGs and each of 28 qualified KO perturbations passing QC (of 30 tested); *PAX6* and *PDX1* are highlighted. e indicates enhancer, HET indicates heterozygous. **c.** Odds ratio for enrichment of the top 500 PerturbScape T2D meta-program genes of each KO (**Methods**) against glucolipotoxicity uDEGs. For perturbations other than *PAX6* and *PDX1*, a single meta-analyzed odds ratio was computed (“Other perturbations”). **d.** Sankey plot of representative shared and condition-specific uDEGs across glucolipotoxicity, *PDX1* KO, and *PAX6* KO, restricted to the top 100 PerturbScape T2D meta-program genes of each KO. **e.** Odds ratio (DEG enrichment, left) and EDC (right) of nutrient-stressor DEGs against db/db (top) and HPAP (bottom) DEGs. **f.** EDC between db/db (top) or HPAP (bottom) DEGs in primary β cells and each KO perturbation in SC-β cells. **g.** T2D heritability enrichment (ENR; left) and standardized effect size (*τ*\*; right) for uDEGs (red) and dDEGs (blue) from the three nutrient-stress programs, HPAP, and db/db DEGs. Throughout, error bars in EDC panels (**b, e, f**) are 95% binomial confidence intervals; error bars in odds-ratio panels (**c, e**) represent exp(log OR ± SE(log OR)); error bars in **g** represent ±1 SE. Asterisks denote significance from a one-sided exact binomial test (EDC > 0.5) in EDC panels, a one-sided Fisher’s exact test in odds-ratio panels, and a one-sided Z-test (ENR > 1 or *τ*\* > 0) in **g**. P-values in **b** and **f** are BH-adjusted; **c**, **e**, and **g** use nominal P-values (*** P < 0.001, ** P < 0.01, * P < 0.05). All analyses were restricted to the β-cell compartment: SC-β cells for the nutrient-stress and CRISPR-perturbation datasets, and primary β cells for the HPAP (human) and db/db (mouse) comparisons. Numerical results are provided in **Supplementary Table S3**.

Having established glucolipotoxicity and *PAX6* KO, *PDX1* KO as convergent perturbations of β-cell identity, we next compared both perturbation classes against transcriptional changes in human T2D islets and the db/db mouse islets. Among nutrient-stress programs, glucolipotoxicity showed the strongest transcriptional concordance with HPAP T2D (OR = 5.1; P= 9e-220, with 1,192 jointly differentially expressed genes; **Figure 4e**), with an effect-direction concordance of 94%. Among genetic perturbations, 13 of 28 KO genotypes showed significant concordance with HPAP T2D (FDR < 5%), with *PAX6* KO again showing the highest alignment (EDC = 0.90) (**Figure 4f**). In contrast, both glucolipotoxicity and KO programs showed weaker concordance with db/db mouse islets (maximum EDC = 0.65 and 0.63, respectively) than with HPAP. This pattern — environmental and genetic perturbations of human β-cells both recapitulating human T2D islets more closely than the prevailing mouse model — establishes that controlled in vitro perturbations in matched human cells provide a more faithful reference for human T2D pathogenesis than animal models that nominally represent the disease at the organism level. Furthermore, glucolipotoxicity and glucotoxicity uDEGs exhibited stronger T2D heritability informativeness than in vivo human and mouse model up- and down-regulated DEGs (uDEGs and dDEGs) (ENR = 4.2–4.4 vs 3.0–3.4; *τ*\* = 0.37 – 0.43 vs 0.09 – 0.28; **Figure 4g**), a difference that persisted after matching the in vivo DEG sets in number to glucolipotoxicity uDEGs (**Supplementary Figure 3**). Strikingly, HPAP T2D-islet DEGs showed no significant heritability informativeness (*τ*\*= 0.09, P= 0.36), likely because the disease-state transcriptome is shaped by a mixture of primary disease mechanisms, compensatory responses, treatment effects, and other secondary consequences of chronic metabolic dysfunction that carry little germline signal. By isolating the primary β-cell response in which genetic variants act, our in vitro programs thus capture germline T2D risk architecture more faithfully than whole-organism human and mouse models.

Together, these findings demonstrate that nutrient stress and genetic loss-of-function perturbations converge on shared transcriptional programs in β cells, with *PAX6*- and *PDX1*-mediated identity networks emerging as central nodes of this convergence, and that controlled in vitro perturbations in human iPSC-derived β-cells more faithfully recapitulate the genetic architecture of human T2D than whole-organism human and mouse models.

### Plasma proteome signatures of β-cell nutrient stress associate with diet and metabolic state

Pancreatic endocrine cells release their products — insulin, glucagon, and stress-responsive proteins — directly into the bloodstream, and we reason that the plasma proteome may carry a measurable projection of β-cell program activity at biobank scale, where direct pancreatic measurement is infeasible. To connect our *in vitro* nutrient-stress signatures to human physiology, we computed individual-level activity scores — stressor Protein Program (sPP) scores — for the three SC-β cell nutrient-stress programs (glucotoxicity, lipotoxicity, and glucolipotoxicity uDEGs), the most T2D heritability enriched class of programs (**Figure 3**), by aggregating normalized abundance of plasma proteins mapping to program genes in 45,956 UK Biobank White British participants with paired Olink Explore proteomics (AUCell method^48^, 2,819 proteins^20^; **Methods**). These sPP scores operationalize the projection step of our “dish-to-biobank” framework, mapping *in vitro* nutrient-induced β-cell stress states onto the plasma proteome. **Figure 5a** summarizes this projection and the downstream phenotypic and genetic analyses.

**Figure 5:**
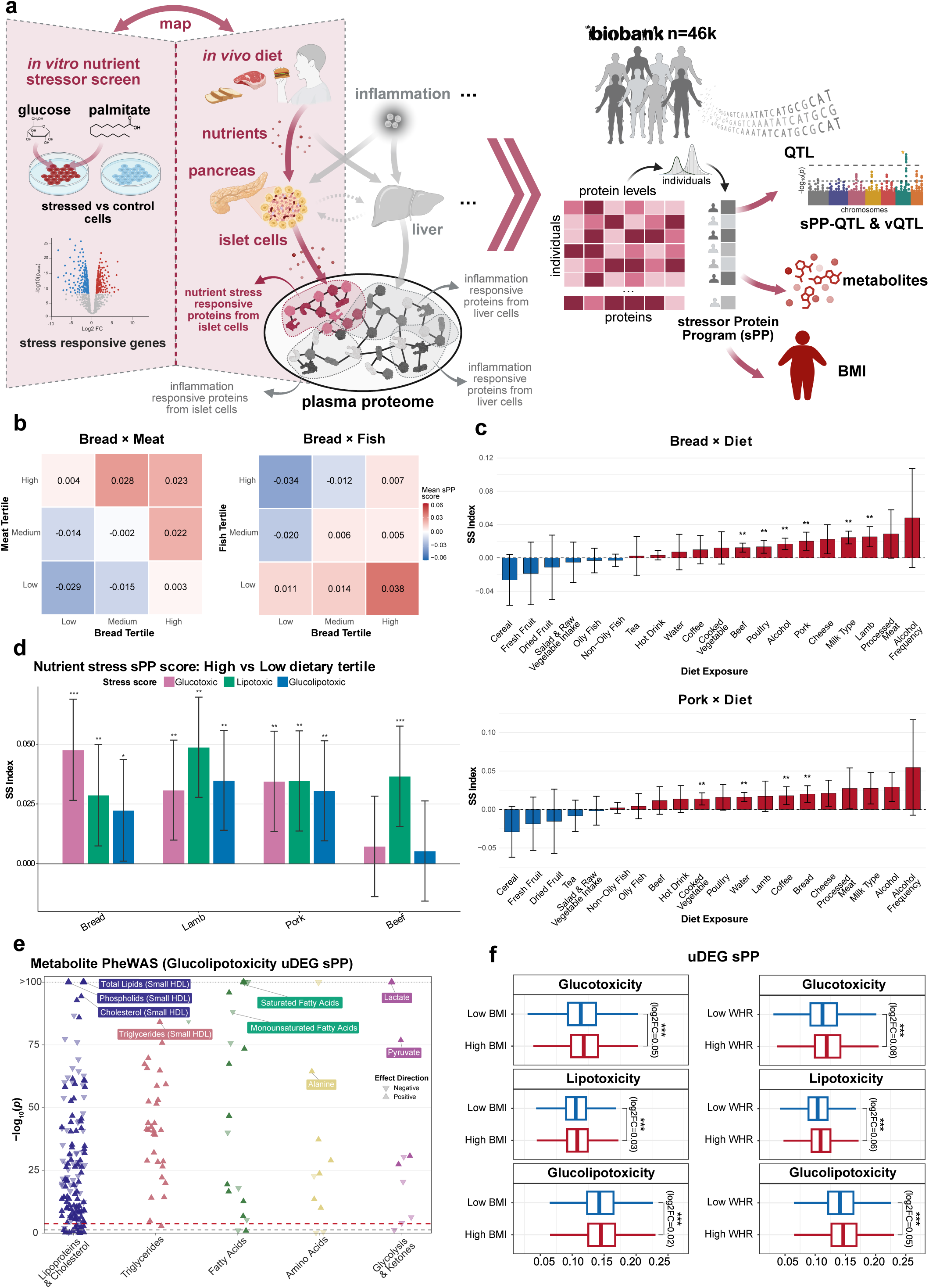
Glucolipotoxicity uDEG sPP captures diet-responsive stress and associates with metabolic traits. **a.** In vitro transcriptomic response of β cells when exposed to glucolipotoxic stress (glucose + palmitate) can be mapped to in vivo secretory plasma protein response from pancreatic islet cells when exposed to nutrient stress associated with proxy glucolipotoxicity diet, comprising of bread and meat intake. The in vitro stress-induced gene programs can be mapped to UKB plasma proteomics (UKB-PPP) using the AUCell method to construct stressor Protein Program (sPP) score per individual. sPP scores are then used for genetic (sPP-QTL and vQTL) analysis, as well as association with metabolites and anthropometric traits. **b.** Mean glucolipotoxicity uDEG sPP across diet tertiles for bread × meat (beef, lamb, pork) and bread × fish (oily and non-oily); color indicates mean sPP. High bread plus high meat intake associates with elevated sPP, with no comparable pattern for fish. **c.** Mean difference in glucolipotoxicity uDEG sPP between individuals in the highest joint tertile of bread × each diet variable (top) or pork × each diet variable (bottom) and the overall mean, across all dietary variables. Red, positive; blue, negative. Asterisks denote P<0.05 based on a permutation-based test on the tertile gradient scores (Methods). Error bars represent 95% CI. **d.** Difference in mean sPP between the highest and lowest dietary tertiles for each of the three nutrient-stress uDEG programs (glucotoxicity, lipotoxicity, glucolipotoxicity) across bread, lamb, pork, and beef. Bread intake associates most strongly with the glucotoxicity uDEG sPP, and lamb and beef with the lipotoxicity uDEG sPP. Error bars represent 95% CI. **e.** PheWAS of glucolipotoxicity uDEG sPP against 251 NMR-quantified plasma metabolites, grouped by metabolite class; point shape indicates effect direction and the dashed line marks the Bonferroni significance threshold. Top associations include small-HDL lipids, saturated fatty acids, lactate, and pyruvate. **f.** Glucotoxicity, lipotoxicity, and glucolipotoxicity uDEG sPP stratified by BMI (low vs high) and waist-to-hip ratio (WHR; low vs high); log₂FC and significance are annotated. Box plots show the median, interquartile range, and whiskers extending to the most extreme observations within 1.5×IQR of the hinges. Asterisks denote significance from one-sided independent samples t-test (*** P < 0.001, ** P < 0.01, * P < 0.05). Numerical results are reported in **Supplementary Table S4**.

We first asked whether sPP scores respond to dietary exposures matching the nutrient-stress conditions used *in vitro* (high glucose, saturated fat). Restricting all analyses to baseline-visit measurements to ensure temporal alignment of proteomic, dietary, and clinical data, we analyzed 20 dietary variables derived from food-frequency questionnaires, encompassing intake frequency of major food groups (**Methods; Supplementary Table S4**). We partitioned each dietary variable into tertiles and assessed sPP scores across both single-diet tertiles and pairwise diet combinations. Given the differences between acute, defined in vitro nutrient exposure and chronic, self-reported dietary variation in humans, concordance between these axes would provide a nontrivial test of the physiological relevance of the sPP scores. Glucolipotoxicity uDEG sPP showed a significant positive association with combined high bread and high red meat intake (meta-analyzed across beef, pork, and lamb; P = 9.2e-03; **Methods**) — a “*proxy glucolipotoxic diet*” of refined carbohydrates plus saturated fat (**Figure 5b**). As a negative control, fish intake (oily and non-oily, rich in unsaturated fatty acids) showed no comparable association with bread (P = 0.51), consistent with saturated fat as the lipid component driving the signal. Bidirectional specificity tests further supported this: holding bread constant, lamb and pork were among the strongest significant associations, while holding red meat constant, bread was among the strongest significant second-component associations for glucolipotoxicity uDEG sPP (**Figure 5c; Supplementary Figure 4a,b, Methods**). This finding supports that carbohydrate x saturated fat combination, rather than either component alone, is the driver of this sPP signature. Importantly, single-diet analyses also recapitulated the mechanistic decomposition we identified *in vitro*: glucotoxicity uDEG sPP showed the strongest association with bread intake, whereas lipotoxicity uDEG sPP showed preferential association with red meat (**Figure 5d, Supplementary Figure 4c**).

To confirm that the sPP scores capture the biochemical signatures of the nutrient-stress states they were designed to measure, we examined associations with circulating metabolites. PheWAS against 251 NMR-quantified circulating metabolites^49,50^revealed strong associations consistent with the glucolipotoxic phenotype. We observed positive associations with lactate (β = 0.17, P = 3e-333), pyruvate (β = 0.08, P = 1.43e-77), and alanine (β = 0.075, P = 3.8e-65) — markers of glycolytic flux and glucose-alanine cycling — and a negative association with glutamine (β = −0.16, P = 1.79e-272), consistent with its consumption as an alternative fuel under metabolic stress (**Figure 5e, Supplementary Table S4**). In parallel, we observed positive associations of glucolipotoxicity uDEG sPP with saturated fatty acids (β = 0.091, P = 1.52e-96), while polyunsaturated and omega-6 fatty acids were negatively associated (β = −0.095, P = 4.56e-106 and β = - 0.095, P = 8.09e-109, respectively). All nutrient-stress uDEG sPP scores showed positive associations with anthropometric markers of metabolic disease, like BMI and waist-to-hip ratio (WHR); individuals in the high BMI and WHR strata showed elevated glucolipotoxicity uDEG sPP relative to low BMI and WHR controls (log₂FC = 0.02 and 0.05, P < 1e-10 in both cases; **Figure 5f**), providing direct phenotypic validation that the in vitro–derived signature captures a disease-relevant physiological state. Together with the dietary and anthropometric associations, these results demonstrate that environmental (diet; external exposure), biochemical (metabolite; internal milieu), and phenotypic (body composition) readouts are consistently aligned with the same underlying β-cell stress-program axis captured by sPP. The top 10 and top 100 most strongly correlated proteins captured 46.6% and 59.6%, respectively, of the proteome-explained variance (R^2^) in glucolipotoxicity uDEG sPP scores, indicating that the signal is distributed across many proteins rather than driven by a small dominant subset **(Supplementary Table S4)**. Finally, we asked whether other tissue-level secretory processes are associated with the pancreatic stress response. Examining the top 50 most-correlated non-program proteins, we found significant tissue-specific expression in liver, adrenal gland, and adipose tissue for several proteins, including *GRHPR* (rank 3), *PEBP1* (rank 15), *LACTB2* (rank 21), and *S100A4* (rank 40) (**Supplementary Figure 4d-g**). These associations indicate that elevated sPP activity co-occurs with liver- and adipose-associated protein signatures, consistent with the score reflecting a whole-body metabolic state rather than a β-cell-restricted signal.

Collectively, these results establish glucolipotoxicity uDEG sPP as a population-scale readout of nutrient-induced β-cell stress programs, showing significant association with dietary, biochemical, and anthropometric phenotypes. This phenotypic anchoring positions sPP scores for the downstream genetic analyses presented in the next section.

### Genetic variants shaping plasma stress-program signatures reveal trans-tissue regulation and disease colocalization

To identify genetic variants influencing circulating stress-program activity, we performed genome-wide association testing of 9,124,756 imputed autosomal variants (MAF ≥ 0.01) against sPP scores in 45,956 White British UK Biobank participants with paired Olink proteomics at the baseline visit (**Methods, Supplementary Table S5, Data Availability**). Genome-wide scanning of the glucolipotoxicity uDEG sPP in SC-β cells — the most disease-informative program from our heritability analyses (**Figure 3**) — revealed 11 trans-acting independent genome-wide significant loci, with prominent signals at *GCKR*, *ARHGEF3*, *SLC22A5,* and *ASGR1/2* (**Figure 6a**). Pathway analysis of genes proximal to these trans-acting loci revealed enrichment for glycerophospholipid metabolism, consistent with metabolic regulation of β-cell nutrient stress (**Figure 6b**). Tissue-expression profiling identified only 2 out of 11 sPP-QTL genes as predominantly liver-restricted (*GCKR* and *ASGR1/2*, **Figure 6c, Supplementary Figure 5a**), while the remaining mapped broadly to other tissues. Because the regulator genes are predominantly non-hepatic, the signal is unlikely to be driven solely by the plasma proteome’s liver-secretion bias^51^, and instead indicates that trans-regulation of the glucolipotoxic stress signature operates through multiple tissues. The two liver-restricted signals, however, provide a focused opportunity to interrogate hepatic regulation of pancreatic stress through the lens of human genetics, as detailed below.

**Figure 6:**
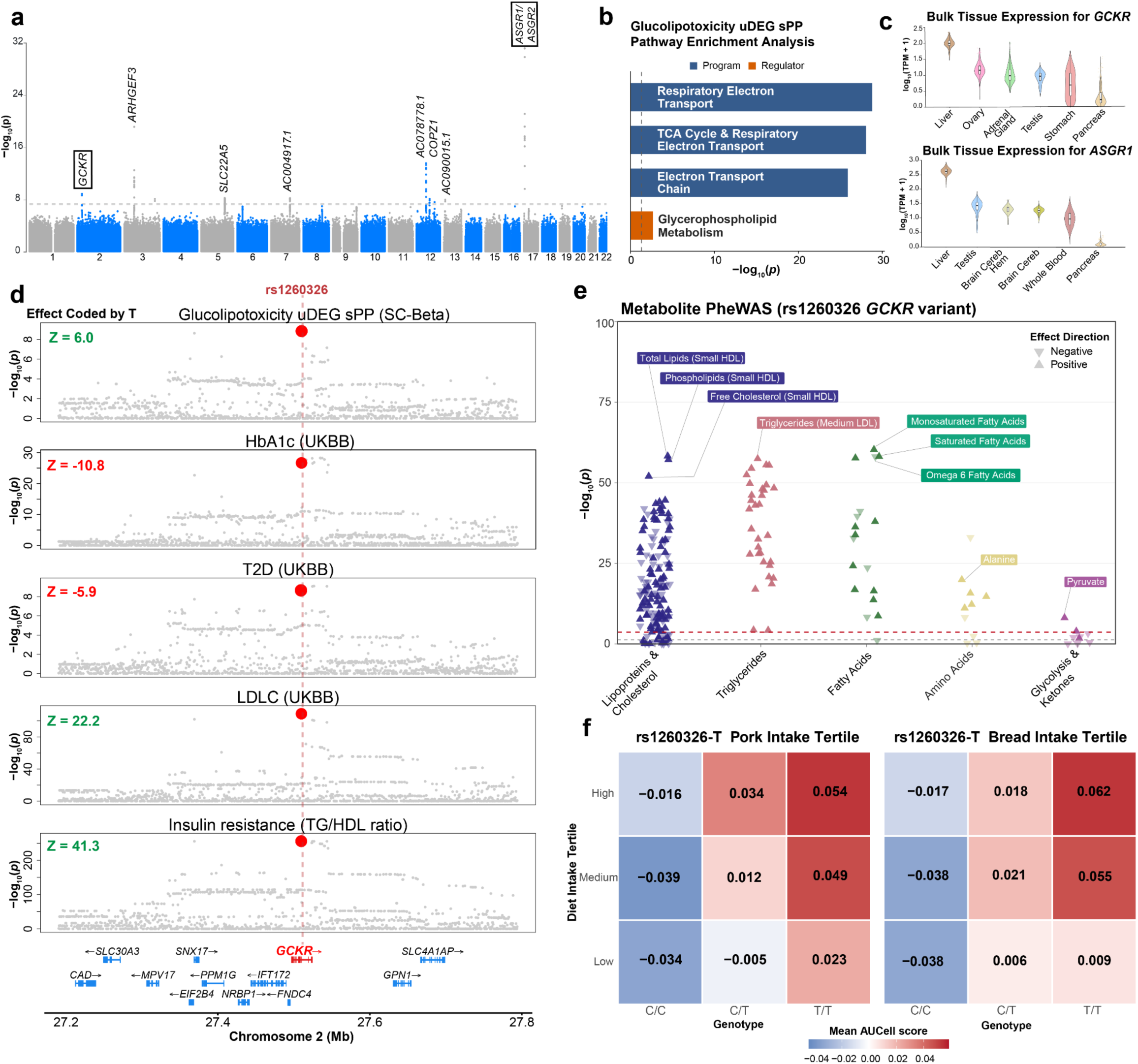
Genetic mapping of the glucolipotoxicity uDEG sPP identifies trans-acting hepatic regulators that colocalize with T2D and lipid genetics. **a.** Manhattan plot of the genome-wide association for the glucolipotoxicity uDEG sPP (N = 45,956) in SC-β cells, identifying 11 independent trans-acting genome-wide-significant loci (P < 5×10⁻⁸; horizontal dotted line). Genes proximal to the sentinel variants are labeled, including the liver-restricted regulators *GCKR* and *ASGR1/2*. **b.** Pathway enrichment (ConsensusPathDB)^99^ for the glucolipotoxicity uDEG sPP program genes (“Program,” blue) and for the trans-acting GWAS regulator genes not represented in the program (“Regulator,” orange). The dashed line marks the significance threshold. **c.** GTEx bulk-tissue expression for *GCKR* (top) and *ASGR1* (bottom) across the highest-ranked tissues; both are most highly expressed in the liver. Neither gene is among the glucolipotoxicity sPP program proteins, identifying them as trans-acting regulators. **d.** Multi-trait colocalization at *rs1260326* (*GCKR* locus, chromosome 2): regional association (−log₁₀P) for the glucolipotoxicity uDEG sPP, HbA1c, T2D, LDL-C, and insulin resistance (TG/HDL ratio), all from UK Biobank. rs1260326-C allele Z-scores are annotated showing opposing glycemic versus lipid effects. **e.** PheWAS of *rs1260326* dosage against 251 NMR-quantified plasma metabolites, grouped by metabolite class; point shape denotes effect direction and the dashed line indicates the significance threshold. Top associations are dominated by lipoprotein, lipid, and fatty-acid species. **f.** Mean glucolipotoxicity uDEG sPP across *rs1260326* genotypes (C/C, C/T, T/T) and pork (left) or bread (right) intake tertiles. Gene-by-environment interaction was significant by an omnibus test for both bread and pork intake (based on an omnibus model; **Methods**). Numerical results are reported in **Supplementary Table S5**.

rs1260326 (P446L), a missense variant in *GCKR*, was the most confidently fine-mapped variant for glucolipotoxicity uDEG sPP (PIP = 0.60) (**Methods**). The variant tags a strong *GCKR* splicing QTL in the liver (GTEx liver, P = 5e-10) but is not a known QTL in the pancreas^52,53^. Although *GCKR* is a known trans-pQTL hotspot^54^, this variant showed stronger association with glucolipotoxicity uDEG sPP compared to size-matched null sPP scores (P=1.34e-09 against null P ranging from 4.76e-07 to 5.66e-01 across 100 null simulations) (**Supplementary Figure 5b**). The biological context for this variant is well established: *GCKR* exerts both inhibitory and stabilizing control over glucokinase; complete loss of *GCKR* reduces GCK protein stability, which can impair net glucokinase activity, whereas the P446L variant weakens *GCKR*-mediated inhibitory control, resulting in increased glucokinase flux^55–57^. *Gckr*-deficient mice develop a diet-conditional metabolic phenotype with diminished hepatic glucokinase activity and impaired glycemic control under high-sucrose/high-fat feeding^55,56^, and P446L knock-in mice on a similar diet recapitulate the pleiotropic glucose-down/lipid-up profile of the human variant association^58^. Our framework extends this in vivo mouse work into human population data through four convergent observations.

First, multi-trait colocalization using ColocBoost^59^ identified strong joint colocalization at P446L (VCP = 0.99) across glucolipotoxicity uDEG sPP-QTL and GWAS for T2D, HbA1c, LDL, and insulin resistance (**Figure 6d**, **Methods**). The T allele increased glucolipotoxicity uDEG sPP, LDL-C and well-established surrogate insulin resistance (TG/HDL-C ratio)^60,61^ markers (Z= 6.0, 22.2 and 41.3 respectively), while lowering HbA1c and T2D risk (Z = −10.8, −5.9 respectively), recapitulating the well-established opposing pleiotropy of P446L on glycemic versus lipid traits^58,62^. Second, metabolite PheWAS of P446L dosage revealed a metabolic signature dominated by lipid-program perturbation: positive associations with lipoprotein particles, triglycerides, and saturated and unsaturated fatty acid species, and a negative association with omega-6 fatty acids, with minimal effects on glycolysis intermediates or ketone bodies (**Figure 6e**, **Methods**). This pattern indicates that the variant’s plasma metabolomic footprint is concentrated in hepatic lipogenic output rather than systemic glycolytic flux, consistent with the LDL-C and TG/HDL-C effects observed at the lipid-trait level. Third, the variant exhibited diet-dependent effects on glucolipotoxicity uDEG sPP: G x pork and G x bread interactions were both significant (omnibus contrast test P = 0.01 and 1.6e-05, respectively; **Methods**, **Figure 6f**), recapitulating at population scale the diet-conditional phenotype originally observed in Gckr-deficient mice^55^. Fourth, we performed genetic mapping of sPP scores corresponding to uDEGs that are glucolipotoxicity-specific (glucolipotoxicity-specific arm), shared between glucolipotoxicity and glucotoxicity uDEGs (glucose arm) and shared between glucolipotoxicity and lipotoxicity uDEGs (lipid arm) (**Supplementary Figure 5c**, **Methods**). This arm-specific pattern recapitulates the finding that the glucose-arm program carries the strongest T2D heritability signal among glucolipotoxicity sub-programs.

The trans-tissue regulatory pattern at *GCKR* is replicated at the second liver-restricted sPP-QTL locus. At *ASGR1/2*, the lead SNP rs186021206 — in linkage with the well-characterized del12 loss-of-function variant^63^ (*r*^2^ = 0.86) shows a directionally concordant pattern with P446L. The G allele at rs186021206 is associated with increased LDL, surrogate insulin resistance, and glucolipotoxicity uDEG sPP (Z= 11.7, 10.9 and 11.5 respectively) (**Supplementary Table S5**). The metabolite PheWAS associations of this variant were also consistent with P446L for lipoproteins and cholesterol metabolites, but weaker for the other metabolites (**Supplementary Figure 5d**). The two loci differ, however, in disease translation: P446L colocalizes robustly with T2D and HbA1c, whereas rs186021206 shows null or minimal effects on both glycemic endpoints (T2D Z= −1.7, HbA1c Z= −2.8). This indicates that a shared trans-tissue effect on plasma lipids and β-cell nutrient stress signatures is not, by itself, sufficient to shift glycemic risk; translation to T2D additionally requires a glucose-lowering action. *GCKR* supplies this through hepatic glucokinase activation, which enhances glucose disposal. However, *ASGR1/2* possibly lacks any comparable glucose-handling effect, so its lipid- and stress-signature associations do not propagate substantially to glycemic endpoints. Together, the *GCKR* and *ASGR1/2* vignettes demonstrate how the dish-to-biobank framework decomposes pleiotropic but directionally opposed effects on glycemic and lipid axes into mechanistic arms of β-cell nutrient stress signatures with diet-dependent modulation.

Genome-wide scanning across all 40 sPP scores spanning nutrient, ER, and inflammatory stress programs identified 497 independent associations (P < 5e-8; *λ*_GC_= 1.005–1.022), with SNP-based heritability (LDSC) reaching significance at 13 of 40 sPPs (heritability Z > 5), including all three nutrient-stress programs (**Supplementary Figure 6a, see Data Availability**). Null sPP scores generated from random size-matched protein sets showed consistently lower heritability z score compared to glucolipotoxicity and glucotoxicity uDEG sPP, but not lipotoxicity uDEG sPP (**Supplementary Figure 6b**), confirming that the genetic signal is specific to biologically coherent stress programs rather than reflecting random protein sets. Among the inflammatory stressors, the cytokine-mix uDEG sPP-QTL was driven by a trans-acting credible set at the *PF4/PPBP/CXCL5* chemokine cluster, with *PF4* and *PPBP* expressed in whole blood and spleen **(Supplementary Figure 6c-e**), implicating immune-tissue chemokine production as a systems-level regulator of β-cell sensitivity to cytokine stress. Multi-phenotype colocalization across the 40 sPP scores and 104 UK Biobank complex traits (**Methods**) identified 87 colocalization events linking sPP-QTLs to disease risk (**Supplementary Figure 6f**, **Supplementary Table S5**).

Together, these findings indicate that the framework’s variant-level resolution generalizes across stressor classes — recovering heritable genetic regulation, trans-tissue biology, and disease-relevant colocalization at population scale.

### Variance-based genetic mapping of glucolipotoxic stress-program signature reveals context-dependent regulation

Standard mean-effect sPP-QTL analysis is limited to variants with uniform effects across individuals; physiologically relevant variants may exhibit context-dependent effects driven by environmental exposures (e.g., dietary patterns) often detectable through variance-based association. Genome-wide variance QTL (vQTL) analysis (OSCA^64^: Bartlett test on RINT-transformed sPP) of glucolipotoxicity uDEG sPP identified rs10774625 — an intronic variant at the pleiotropic 12q24 locus encompassing *ATXN2* and *SH2B3* — at suggestive significance (P = 9e-07; **Figure 7a**, **Supplementary Figure 7a, Methods**). Similar results were observed using the Levene test instead of Bartlett (P=4.2e-06). vQTL signals predominantly reflect gene-environment interactions, and such interactions are typically underpowered to detect at population scale relative to main genetic effects^65,66^. This variant was not detected in standard mean-effect sPP-QTL analysis (P = 0.21; **Figure 7a**), indicating that its influence on glucolipotoxicity uDEG sPP operates through variance heterogeneity across genotypes rather than mean differences — a signature of effect modification by unmeasured contextual factors. The 12q24 locus is associated with an exceptionally broad phenotypic spectrum including T1D, T2D, blood cell traits, and cardiovascular disease (**Supplementary Table S7**), consistent with context-dependent pleiotropy at a single regulatory element.

**Figure 7:**
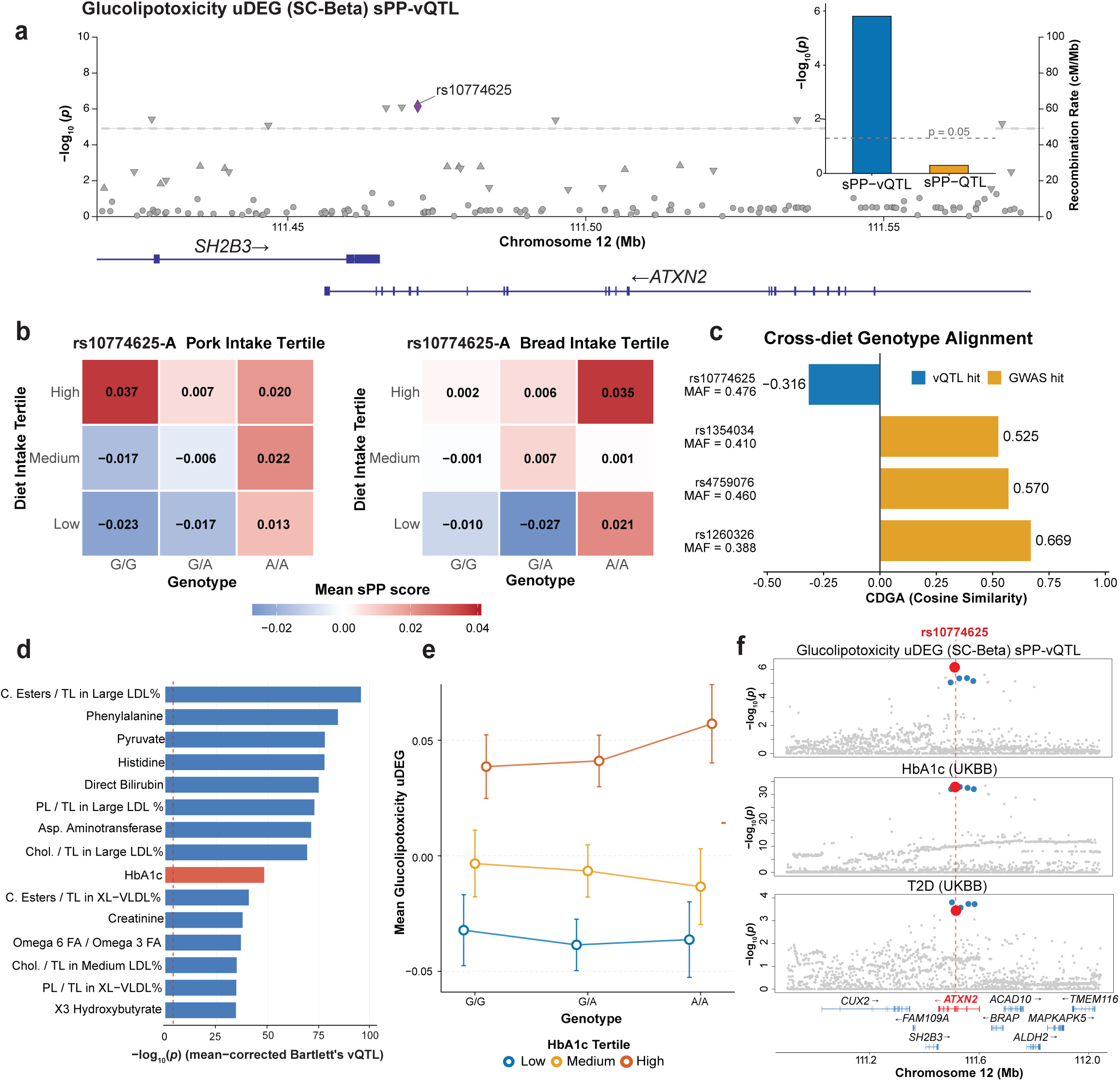
rs10774625 is a variance QTL for glucolipotoxicity sPP with metabolic and disease relevance. **a.** Variance-QTL (vQTL) mapping (Bartlett’s test on RINT-transformed scores) of the glucolipotoxicity uDEG sPP score in SC-β cells across the 12q24 *ATXN2/SH2B3* locus; *rs10774625* reaches suggestive significance (P = 9×10⁻⁷). Horizontal line, suggestive threshold; right axis, recombination rate; gene models below. Inset: −log₁₀P for rs10774625 from sPP-vQTL versus mean-effect sPP-QTL mapping, showing a variance-specific effect (mean-QTL P = 0.21). **b.** Mean glucolipotoxicity uDEG sPP across rs10774625 genotypes (G/G, G/A, A/A) and pork (left) or bread (right) intake tertiles; color indicates mean sPP. The genotype effect diverges in opposite directions between the two diets at the highest tertile. **c.** Cross-diet genotype alignment (CDGA): the cosine similarity between a variant’s dosage-effect vectors in bread- versus meat-intake contexts, where negative values denote opposing (discordant) effects. The vQTL-discovered *rs10774625* is discordant (CDGA = −0.32), whereas mean-effect sPP-QTLs (*rs1260326* at *GCKR*, *rs1354034*, *rs4759076*) are concordant (0.52–0.67). MAF is shown for each variant. **d.** Traits with the strongest variance association with *rs10774625* dosage, ranked by −log₁₀P (Bartlett’s vQTL), across 251 NMR metabolites and 33 blood-biochemistry biomarkers; cholesteryl esters in large LDL is the top association and HbA1c ranks 9th (highlighted). **e.** Mean glucolipotoxicity uDEG sPP across rs10774625 genotypes (G/G, G/A, A/A), stratified by HbA1c tertile (low, medium, high); the dosage effect is evident only in the high-HbA1c stratum. Error bars denote 95% CI. **f.** Multi-trait colocalization at *rs10774625* (*12q24*): regional association (−log₁₀P) for the glucolipotoxicity uDEG sPP-vQTL (top), HbA1c (middle), and T2D GWAS (bottom, indicating a shared signal across the *ATXN2/SH2B3* locus (VCP = 0.61). Numerical results are reported in **Supplementary Table S6**.

The variant’s diet-dependent effects supported the vQTL association, also clarifying why standard mean-effect mapping failed to detect it. Notably, we observed opposing dietary modulation of the rs10774625-A allele: at high bread intake, the variant’s effect on glucolipotoxicity uDEG sPP increased monotonically with A allele dosage, whereas at high pork intake the strongest sPP elevation occurred at the G/G genotype (**Figure 7b**). Consistent with a vQTL, neither G x bread nor G x pork was individually significant (P = 0.31, 0.30): such single-exposure tests are underpowered, and the opposing modulation across these dietary exposures cancels the marginal mean effect while inflating variance. To quantify the opposing modulation, we computed cross-diet genotype alignment (CDGA), the cosine similarity between a variant’s dosage-effect vectors estimated separately in bread- and meat-intake contexts (**Methods**), with negative values denoting opposing effects. As diet entered neither the in vitro–derived sPP score nor the variance test, this cross-diet structure is an independent in vivo observation. rs10774625 showed negative alignment (CDGA = −0.32) in contrast to other fine-mapped mean-effect sPP-QTLs with sufficient homozygous representation (>1,000 carriers); rs1260326 at *GCKR*, rs1354034, and rs4759076 all showed concordant cross-diet effects (CDGA = 0.52-0.67) (**Figure 7c)**. These opposing diet-specific effects cancel in the population mean, accounting for the null mean-effect association, while persisting as the variance heterogeneity that vQTL detects.

To extend this contextual characterization beyond diet, we performed vQTL scans of the rs10774625 variant dosage across 251 metabolites and 33 blood biochemistry biomarkers in UK Biobank (**Methods, Supplementary Table S6**). We identified 59 traits with significant variance association at rs10774625 (P < 1e-05), with cholesteryl esters as the strongest variance association (rank 1) and HbA1c also among the top hits (rank 9; **Figure 7d, Supplementary Figure 7b)**. We focused on HbA1c as a clinically interpretable axis of chronic glycemic state. Genotype-stratified analysis of glucolipotoxicity uDEG sPP across HbA1c tertiles revealed that homozygous minor-allele carriers exhibited elevated sPP specifically when HbA1c was high, with minimal genotype effect at low and medium HbA1c levels (**Figure 7e**), consistent with glycemic state modifying the variant’s effect on the stress signature. As HbA1c is itself shaped by glycemic regulation, this stratification is descriptive and does not establish causal direction. Multi-trait colocalization using ColocBoost^59^ at rs10774625 identified joint colocalization (VCP = 0.61) between glucolipotoxicity uDEG sPP-vQTL, HbA1c GWAS, and T2D GWAS (**Figure 7f**), indicating a shared genetic signal across these phenotypes (VCP is analogous to fine-mapping PIP and a score of > 0.5 is considered very strong). Mean-and variance-based mapping of stressor-program scores thus capture complementary axes of T2D genetic risk at this locus.

Our findings extend the *ATXN2* phenotypic spectrum to context-dependent regulation of β-cell nutrient stress signature in T2D and show that variance-based genetic mapping can recover disease-relevant variants that mean-effect analysis can miss.

## Discussion

Here, we establish a generalizable framework linking controlled in vitro perturbation biology to human disease genetics and physiology. Rather than asking whether an in vitro model perfectly recreates the in vivo conditions, this framework asks a more testable question: whether programs induced by defined perturbations align with the genetic, dietary, metabolomic, and disease structure observed in humans at scale. By integrating single-cell stress responses in human stem cell–derived islets with population-scale proteomics and genetic analyses, we map defined nutrient exposures (glucose and palmitate) to transcriptional programs, circulating protein signatures, and human genetic variation (**Figure 1**). Stem cell–derived islets align more closely in their stress response with human genetic perturbation data and human T2D islet transcriptomes than the db/db mouse model does, helping to bridge a longstanding gap between experimental models and human pathophysiology.

Applied to T2D, this framework identifies glucolipotoxicity as a genetically anchored model of β-cell dysfunction: a coordinated state of insulin induction, ER and mitochondrial stress, and eroding β-cell identity, consistent with a compensatory response that may progress to β-cell failure under sustained stress^67,68^. Among its three arms — glucose-driven, lipid-driven, and glucolipotoxicity-specific — the glucose-driven arm carries the strongest T2D heritability and is also the arm preferentially modulated by the *GCKR* variant, positioning it as the principal axis linking the program to disease risk. Genetic disruption of *PAX6* and *PDX1* produces a transcriptional state aligning with the glucolipotoxic program, indicating that environmental and genetic perturbations converge on a shared phenotype. Finally, monogenic, rare-variant, and common-variant signals — despite minimal mutual overlap — each converge on this program, pointing to a core of β-cell dysfunction that spans the allele-frequency spectrum.

Projection of stress-induced programs onto the plasma proteome shows that the nutrient-stress signature is measurable in human populations and shaped by both dietary exposures and genetic variation. These circulating signatures reflect not only β-cell–intrinsic processes but also cross-organ metabolic interactions, as illustrated by trans-acting loci such as *GCKR, ASGR1/2,* and *ATXN2*. Importantly, a subset of these loci exhibit diet-dependent effects, indicating that gene-by-environment interactions analogous to those in our in vitro factorial design are detectable in vivo at population scale. As an RNA-binding protein involved in translational control and stress responses, *ATXN2* also illustrates the broader translational relevance of these loci: antisense oligonucleotide therapeutics targeting it have already shown promise in preclinical models^69,70^.

These findings open several avenues for translational and mechanistic discovery. First, the plasma-as-a-bridge strategy provides a principled route to dissect inter-organ signaling by identifying trans-acting regulators (e.g., liver-derived factors) that modulate β-cell stress programs. Second, stressor Protein Programs (sPPs) offer candidate biomarkers for early detection and risk stratification, as well as a substrate for testing gene-by-environment interactions in population cohorts, enabled by the direct correspondence between our in vitro stressors (glucose and palmitate) and their in vivo dietary analogs (refined carbohydrate and saturated fat intake). Third, the alignment between specific genetic perturbations and stress responses — exemplified here by *PAX6* and *PDX1* under glucolipotoxicity — motivates combinatorial perturbation x stress designs to directly dissect how genetic and environmental insults interact to destabilize β-cell identity. Fourth, genes shared across stress-induced programs, genetic perturbation programs, and T2D common-and rare-variant associations represent high-confidence candidates for therapeutic target development.

This study establishes a controlled dish-to-biobank framework, and several extensions could broaden its generalizability and mechanistic resolution. The in vitro perturbations were performed in a single iPSC line, providing a controlled but necessarily narrow view of inter-individual variability; iPSC “village” designs^71,72^ that pool multiple donor lines under shared perturbations represent a natural next step, allowing genetic background to be tested as a modulator of stress program intensity and providing a “cohort in a dish” that bridges single-donor mechanism to population-scale variability. Although stem cell–derived islets capture core β-cell biology, their developmental immaturity relative to adult primary islets^73^ means some aspects of the stress response may differ from in vivo β-cells. Primary human islets provide an important adult-tissue benchmark, but donor availability, variable islet quality, isolation and culture stress, and batch effects make controlled factorial perturbation designs challenging; future studies that compare SC-islets and primary islets across matched stressors and timepoints will help define which stress programs are model-invariant versus context-specific. The perturbation panel itself can also be expanded — moving from the small set of β-cell identity regulators perturbed here toward broader disease-gene libraries to systematically map which gene–environment combinations converge on which disease-relevant cellular programs. The population-scale proteomic, metabolomic, dietary and physiological trait analyses were restricted to UK Biobank participants of European ancestry; extension to All of Us^74^ and Biobank Japan^75^ will be essential as comparable proteomic panels mature in those cohorts, and analogous designs across more diverse populations will refine which trans-tissue regulators are shared and which are population-specific. In addition, dietary exposures and plasma molecular measurements in UK Biobank are observational and largely cross-sectional; questionnaire-derived diet variables are imperfect proxies for long-term nutrient exposure, and a single plasma measurement may not capture chronic metabolic state. Therefore, diet-sPP associations are not individual-level causal estimates of diet, but rather a population-scale concordance assessment of whether sPPs vary along environmental and metabolic axes in humans.

Most consequentially, the dish-to-biobank framework is naturally applicable to any cell type with measurable secretory contributions to the plasma proteome — making liver, kidney, immune, and adipose disease contexts immediate extensions. For tissues with limited circulating representation, such as brain neurons, alternative population-scale molecular readouts (for example, cerebrospinal fluid proteomics for neurological diseases) provide an analogous bridge between controlled cellular perturbation and human disease genetics. This framework thus offers a generalizable template for converting in vitro mechanism into population-scale insight, with each axis of expansion — donor diversity, perturbation breadth, tissue and disease scope — opening a distinct avenue for the field.

## Data Availability

The single-cell RNA-sequencing data of nutrient-stressed SC-islets generated in this study, including the raw reads and processed count matrices, have been deposited in the Gene Expression Omnibus (GEO) under accession <u>GSE333721</u>. Differential-expression results and stress-program (sPP) gene sets are provided in **Supplementary Tables S1** and **S4**, and at https://zenodo.org/records/20514606.

Summary statistics for the sPP-QTL and vQTL analyses have been deposited in Zenodo (https://zenodo.org/records/20514606) and will be uploaded to the NHGRI-EBI GWAS Catalog upon publication. HbA1c burden test summary results are also available in the same Zenodo link. The summary statistics for UK Biobank used in this paper are available at https://data.broadinstitute.org/alkesgroup/UKBB. the surrogate insulin resistance GWAS has been fetched from GWAS catalog (https://www.ebi.ac.uk/gwas/publications/38200128). Individual-level UK Biobank data (Olink Explore proteomics, NMR metabolomics, dietary, anthropometric, and genetic data) are available to approved researchers through the UK Biobank Access Management System (https://www.ukbiobank.ac.uk) through the Research Analysis Platform (RAP); analyses in this study were performed under Application 16549. These data cannot be redistributed by the authors under the terms of UK Biobank access.

This study used the following publicly available datasets: HPAP (Human Pancreas Analysis Program; https://hpap.pmacs.upenn.edu; ref^25^); the db/db mouse islet dataset (GSE211799; ref^24^); the SC-islet CRISPR Perturb-seq dataset (GSE313516; ref^27^); the ER/inflammatory stress single-cell screen (GSE237448; ref^18^); GTEx v8 (https://gtexportal.org) and 1000 Genomes Project Phase 3 (https://www.internationalgenome.org).

## Code Availability

Single-cell processing, differential-expression, sPP scoring using AUCell, omnibus test, CDGA calculation: https://github.com/Deylab999MSKCC/glucolipotoxicity

sc-linker: https://github.com/kkdey/GSSG, https://github.com/karthikj89/scgenetics LDSC software: https://github.com/bulik/ldsc/wiki

REGENIE software for performing sPP-QTL, also HbA1c burden WES: https://rgcgithub.github.io/regenie/ Large scale genetic analysis using the DNAnexus platform: https://platform.dnanexus.com/

ENCODE-rE2G enhancer-gene linking: https://www.encodeproject.org/software/distal-regulation-encode_re2g/

Cell Ranger (single-cell transcriptome processing) — https://www.10xgenomics.com/support/software/cell-ranger

Seurat (clustering, FindMarkers DE, cell-cycle scoring) — https://satijalab.org/seurat/

SuSiE / susieR (fine-mapping, PIP > 0.2**) —** https://github.com/stephenslab/susieR OSCA (vQTL test) - https://yanglab.westlake.edu.cn/software/osca/

ColocBoost (multi-phenotype colocalization) - https://github.com/StatFunGen/colocboost Perturbscape method (perturbation program atlas and meta-program calling): https://github.com/Deylab999MSKCC/perturbscape

## Acknowledgement

We would like to thank all the members of Dey laboratory and Huangfu laboratory. We acknowledge the assistance from the following Memorial Sloan Kettering Cancer Center Cores: Flow Cytometry and Integrated Genomics Operation. This study was funded in part by National Institutes of Health (NIH) grants R01HG014008 (K.K.D.), U01HG012051 and UM1HG012654 (D.H.), and NIH/NCI MSKCC Cancer Center Support Grant P30CA008748. This research has been conducted using the UK Biobank Resource under Application Number 16549.

## Author contributions

X.W., D.H., and K.K.D. designed experiments. X.W., H.L., A.L., B.T., D.H., and K.K.D. interpreted results. X.W. performed the experiments. H.L. performed disease heritability analyses and comparisons between stress and perturbation-induced signatures. B.T. performed DEG analysis of scRNA-seq data from stress, mouse models and human T2D islets. A.L. and P.S.G. performed sPP scoring and subsequent phenome-level (diet, metabolite) and genetic analyses. D.L. generated the CRISPR-knockout scRNA-seq dataset. N.Z. assisted with the scRNA-seq experimental set up. B.W. assisted with optimization of glucolipotoxicity treatment conditions in SC-islets. T.A.A. and A.L. contributed to the genetic pipeline development. X.C. performed fine-mapping and colocalization. N.H. and B.T. performed scRNA-seq alignment, quality control, and pre-processing. C.A.L., G.W. and K.K.D. supervised genetic analyses. X.W., D.H., and K.K.D. wrote the manuscript.

## Methods

### Cell culture system and maintenance

XM001 hiPSCs were maintained in Essential 8 (E8) medium (Thermo Fisher Scientific, A1517001) on vitronectin (Thermo Fisher Scientific, A14700) pre-coated plates at 37°C with 5% CO_2_. For regular maintenance, cells were passaged every 3-4 days using Accutase (STEMCELL Technologies, 07920) for dissociation. The Rho-associated protein kinase (ROCK) inhibitor Y-27632 (10 μM; Selleck Chemicals, S1049) was added to the E8 medium for the first 24 h after passaging or thawing of hiPSCs. Cells were regularly tested and confirmed to be mycoplasma free by the Memorial Sloan Kettering Cancer Center (MSKCC) Antibody & Bioresource Core Facility.

### Stem cell-derived islet differentiation

hiPSCs were seeded at a density of 0.75 ×10^5^ cells/cm^2^ on Matrigel (Corning, 354253) pre-coated plates in E8 medium with 10 μM Y-27632. After 48 hours, cells were washed with DPBS once and differentiated following previously described protocols with some modifications^76,77^. During stage 1 to stage 6 day 7, cells were cultured in 6-well plates. On stage 6 day 7, cells were dissociated with Accutase. 5 ×10^6^ cells were resuspended in 5 ml ESFM medium and transferred into a single well of ultra-low attachment 6-well plates (Corning, 3471). The plate was incubated on an orbital shaker at 100 rpm.

**Stage 1 (3 d):** S1/2 medium supplemented with 100 ng/ml Activin A (Bon Opus Biosciences, C687-1MG) and 3 μM CHIR99021 (Stemgent, 04-0004-10) for day 1. S1/2 medium supplemented with 100 ng/ml Activin A and 0.3 uM CHIR99021 for day 2. S1/2 medium supplemented with 100 ng/ml Activin A for day 3. S1/2 base medium: 500 ml MCDB 131 (Life Technologies, 10372019) supplemented with 2.07 ml 45% glucose (MilliporeSigma, G7528), 0.75 g sodium bicarbonate (MilliporeSigma, S5761), 2.59 g BSA (Proliant, 68700), 5.1 ml GlutaMAX (Invitrogen, 35050079).

**Stage 2 (2 d)**: S1/2 medium supplemented with 50 ng/ml KGF (PeproTech, AF-100-19) and 0.25 mM vitamin C (VitC) (Sigma-Aldrich, A4544).

**Stage 3 (2 d):** S3/4 medium supplemented with 50 ng/ml KGF, 0.25 mM VitC, 0.25 uM SANT-1 (Sigma, S4572), 2 μM retinoic acid (MilliporeSigma, R2625), 200 nM LDN (Stemgent, 04-0019) and 500 nM TPPB (TOCRIS, 5343). S3/4 base medium: 500 ml MCDB 131 supplemented with 0.52 ml 45% glucose, 0.877 g sodium bicarbonate, 10 g BSA, 2.5 ml ITS-X (Life Technologies, 51500056), 5.2 ml GlutaMAX.

**Stage 4 (4 d)**: S3/4 medium supplemented with 50 ng/ ml KGF, 0.25 mM VitC, 0.25 μM SANT-1, 0.1 μM retinoic acid, 200 nM LDN and 500 nM TPPB.

**Stage 5 (7 d):** S5 medium supplemented with 10 µM ALK5i II (Cayman Chemical Company, 14794-5), 0.25 mM VitC, 0.25 μM SANT-1, 0.1 μM retinoic acid, 1 µM T3 (Sigma-Aldrich, T6397), 20 ng/ml Betacellulin (R&D, 261-CE-050/CF), 10 μg/ml Heparin (Sigma-Aldrich, H3149) and 1 µM γ-Secretase Inhibitor XXI (EMD Millipore, 565790). S5 base medium: 500 ml MCDB 131 supplemented with 4 ml 45% glucose, 0.877 g sodium bicarbonate, 10 g BSA, 2.5 ml ITS-X, 5.2 ml GlutaMAX.

**Stage 6 (14 d):** ESFM medium. ESFM medium: 500 ml MCDB 131 supplemented with 0.52 ml 45% glucose, 10.5 g BSA, 5.2 ml GlutaMAX, 5.2 ml NEAA (Invitrogen, 11140050), 1 μM ZnSO4 (MilliporeSigma, 108883), 523 μl Trace Elements A (Corning, 25-021-CI), 523 μl Trace Elements B (Corning, 25-022-CI), 10 ug /ml Heparin.

### Nutrient stress treatment of SC-islet cells

A single differentiation of XM001 was performed; at stage 6 day 14 the resulting SC-islet pool was divided into six aliquots, each assigned to one nutrient-stress condition and treated independently for 72 h. The four primary nutrient stress conditions were (i) 8 mM glucose + BSA vehicle control (Cayman Chemical, 29556), (ii) 8 mM glucose + 1 mM BSA-conjugated palmitate (Cayman Chemical, 29558), (iii) 33 mM glucose + BSA vehicle control, and (iv) 33 mM glucose + 1 mM BSA-conjugated palmitate. Additionally, we included two secondary stress experiments comprising 8 mM glucose (no BSA) and 33 mM glucose (no BSA). After 72 h of treatment, SC-islets were dissociated into single cells, collected, and cryopreserved in Bambanker cell freezing medium (Fujifilm, 302-14681).

This isogenic design eliminates donor-specific genetic modifiers as a confound, ensuring that transcriptional differences between conditions are attributable to the stressor rather than to genetic heterogeneity between lines — the same principle that underlies other isogenic CRISPR perturbation designs in human pluripotent stem cells^78,79^. This design also enabled the high per-condition cell numbers (5,067-7,686 after QC) required to resolve stressor-specific DEG programs at single-cell resolution, which would not have been feasible in a multi-donor village culture design, where per-condition cell counts are diluted across genetic backgrounds. Biological validity of the resulting stress-response genes is established through orthogonal evidence independent of line identity – concordance with human T2D islet transcriptomes (HPAP), db/db mouse model signatures, CRISPR KO perturbation alignment (*PAX6*, *PDX1*), rare-variant burden convergence, MODY gene set overlap, and common-variant heritability enrichment.

### Single-cell RNA library preparation, sequencing and analysis

On the day of scRNA-seq library preparation, frozen cells were thawed at 37°C, resuspended in PBS containing 2% BSA and then incubated with Live/Dead Fixable Violet Dead cell stain (Invitrogen, L34955) in FACS buffer for 15 minutes at room temperature. Live cells were sorted by FACS and loaded onto the Chromium Controller with a targeted collection of 10,000 cells per reaction (10x Genomics Chromium Single Cell 3′ Reagent Kit v3.1 User Guide). cDNA and sequencing libraries were prepared according to the manufacturer’s instructions and sequenced on NovaSeq 6000 platform.

Transcriptome libraries were processed with Cell Ranger v8.0.0 using the GRCh38-2024-A reference and default parameters, generating gene-by-cell count matrices. Downstream analyses were performed in Seurat v.5.0.3^80^. We applied a two-stage quality control strategy designed to balance broad cluster representation with precise cell quantification for downstream differential expression. In the first stage, we applied relaxed filters (removing only cells with fewer than 200 detected genes or more than 50% mitochondrial transcripts) and performed Louvain clustering at resolution 0.6 on the resulting cells (N = 42,053 cells; 389,403,101 UMIs). Clusters were annotated based on marker gene profiles, identifying 8 transcriptionally distinct cell populations. In the second stage, we applied stringent QC filters to the annotated dataset, retaining cells with more than 1,000 and fewer than 7,500 detected genes (nFeature_RNA) and less than 25% mitochondrial transcripts, while preserving the original cell-type annotations (N = 38,350 cells; 382,245,713 UMIs retained). This two-stage ordering reflects the biological principle that cell types differ in their native distributions of gene detection and mitochondrial expression, and is supported by previous research^81–83^. Applying stringent filters only after cluster annotation ensures that downstream differential expression operates on high-quality cells while clustering captures the full cell-type landscape.

### Identifying differentially expressed genes (DEGs) from nutrient stress scRNA data

For the nutrient stress experiments, differential expression (DE) testing was performed using the Wilcoxon rank sum test to identify genes with significant expression changes between each stressor condition and baseline, within each annotated cell type. This test treats each cell as an independent observation of stressor response given a fixed genetic background. Genes were retained if they exhibited a minimum effect size (|log₂FC| > 0.1) and sufficient expression (detected in >1% of cells in at least one condition). Bonferroni correction was applied based on the total number of genes, and genes with adjusted P < 0.05 were classified as significantly differentially expressed genes (DEGs), further split into up-regulated (uDEG; log₂FC > 0) or down-regulated (dDEG; log₂FC < 0) relative to baseline. This DE analysis strategy is consistent with the default implementation of the FindMarkers() function in the Seurat R package^80^. Additionally, we performed variance component analysis using the variancePartition BioC package^84^. We applied the same DE protocol to identify cell type-specific DEGs in two additional contexts: (i) db/db islets compared to matched wild-type controls^24,85,86^, and (ii) each of knockout genotype compared to unperturbed cells at SC-β cell stage of the time-resolved CRISPR-KO single cell RNA-seq data^27^. For the single cell RNA-seq data, the stringent adjusted P < 0.05 threshold yielded few DEGs per perturbation (<100 genes for 12 perturbations), reflecting limited statistical power due to the single-cell resolution of individual perturbations (CRISPR-KO: 1-464 cells per condition, mean = 70.56). To enable comparative analysis with nutrient stress signatures, we therefore used a nominal P < 0.05 threshold for these perturbation datasets. For ER and inflammatory stress conditions, DEGs were obtained directly from the original publication^18^, as complete cell-level metadata required for uniform reprocessing were not available. To identify genes differentially expressed in T2D β cells, we analyzed publicly available single-cell RNA-sequencing (scRNA-seq) datasets from the Human Pancreas Analysis Program^87^ (HPAP; https://hpap.pmacs.upenn.edu/). Differentially expressed genes (DEGs) were identified by comparing β cells from T2D donors with β cells from non-diabetic control donors using the FindMarkers() function in Seurat.

### Sc-linker disease heritability analysis of DEGs

We used sc-linker^4^ to quantify disease heritability enrichment for gene programs derived from differential expression analysis. For each stressor or perturbation condition, up-regulated and down-regulated DEGs (uDEGs and dDEGs) in a cell type were treated as separate gene sets. SNP-to-gene linking was performed using ENCODE-rE2G^35^ (union across bio-samples), which has been benchmarked to outperform the ABC and Roadmap-based linking strategies employed in the original sc-linker implementation. For each gene set G, we generated a binary 0/1 annotation for 9,991,229 variants with minor allele count ≥5 from the 1000 Genomes Project European samples^88^, based on whether the variant was linked to a gene in the program. Disease heritability analysis for diabetes-related traits (average N=209,644) (**Supplementary Table S2**) was performed using stratified LD score regression (S-LDSC)^89,90^, conditioning on 97 baseline-LD (v2.2) annotations comprising coding, conserved, and LD-related features^36,37^. We report two metrics from sc-linker: (i) heritability enrichment and (ii) standardized effect size (*τ*^∗^). Heritability enrichment is defined as %(*h*^2^(*G*))/%*SNP*(*G*) for gene set *G* – where %(*h*^2^(*G*)) corresponds to the fraction of heritability captured by variants linked to genes in the gene set and %*SNP*(*G*) corresponds to the fraction of variants out of all variants that are linked to the genes in the program. Standardized effect size (*τ*^∗^) is defined as the proportionate change in per-variant heritability associated with a one standard deviation increase in the value of the sc-linker annotation, conditional on other annotations included in the model. *τ*^∗^ is of the form

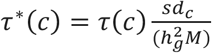

where *τ*(*c*) is the contribution of annotation *c* to the per-SNP heritability of a disease conditional on other annotations. The per-SNP heritability is modelled as 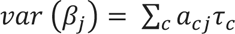, where *a_cj_* denotes the value of annotation *c* in each SNP *j*. *sd_c_*, is the standard error of annotation *c*, 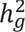 is the total SNP heritability, and *M* is the total number of commonly varying SNPs on which this heritability is computed (equal to 5,961,159 in our analyses).

### PerturbScape meta-program analysis

PerturbScape is a framework that analyzes how genetic perturbations relate to disease risk. It first extracts gene programs from CRISPR Perturb-seq experimental data using multiple contrastive and non-contrastive embedding methods for each perturbation, then evaluates how strongly each program is associated with disease genetic risk. Significant programs are combined into a single “meta-program”, which assigns a score for each gene reflecting its contribution to the perturbation’s disease-relevant downstream effects by training on the disease GWAS score using the MAGMA method^39^; this is analogous to generating PoPS score^91^ but by restricting the features to a perturbation-specific program atlas. In our analysis, the top 500 genes from each perturbation’s meta-program for T2D were used to assess their overlap with glucolipotoxicity uDEGs. Details of PerturbScape are provided in the companion paper (Turhan*, Lee* in preparation, see **Code Availability**).

### Curated set of diabetes related gene sets

The 175 expertly curated gene list associated with monogenic and polygenic forms of diabetes were generated by combining multiple data resources, including (i) monogenic diabetes genes from PhenoApp shared by Dr. Elisa De Franco, (ii) monogenic diabetes genes from recent Bonnefond et al^92^ paper, (iii) curated T2D effector gene predictions from the Common Metabolic Diseases Knowledge Portal (see **Data Availability**), (iv) MODY genes from the recent Bonnefond et al^92^ paper, (v) neonatal diabetes risk genes from Bonnefond et al., and (vi) syndromic genes from Bonnefond et al. Second, outside of these expert-curated genes, we considered a broader 561 MODY-associated gene set from ref^44^.

Rare-variant, gene-based burden testing for glycated hemoglobin (HbA1c) was performed on UK Biobank whole-exome sequencing (WES) data using REGENIE (v4.1.2). In step 1, a whole-genome regression model was fit on directly genotyped array variants with leave-one-chromosome-out (LOCO) predictions to account for relatedness and population structure; in step 2, qualifying variants were collapsed into gene-based masks. Variants were annotated and aggregated into the seven recommended masks defined by predicted consequence and missense deleteriousness (LoF, synonymous, mis-sense, LoF+mis-sense, LoF+synonymous, mis-sense+synonymous, LoF+mis-sense+synonymous). Only variants with MAF < 0.001 (0.1%) were retained; restricting to this threshold limits contamination from lower-frequency variants in linkage disequilibrium with common-variant disease signals^93^. All models were adjusted for age, sex, age^2^, age × sex, the first 10 genetic principal components, and genotyping array. Burden summary statistics are made publicly available (see **Data Availability**).

### UK Biobank cohort and phenotype data

UK Biobank (UKB) is a prospective, population-based cohort of approximately 500,000 individuals aged 40–69 years at recruitment (2006–2010)^19,20^. Participant data include genome-wide genotyping, exome sequencing, whole-body magnetic resonance imaging, electronic health record linkage, blood and urine biomarkers, and physical and anthropometric measurements. Proteomic data were generated by the UK Biobank Pharma Proteomics Project (UKB-PPP), a consortium of 13 biopharmaceutical companies. We analyzed plasma protein measurements from 45,956 White British individuals at the baseline assessment (2006–2010). Normalized Protein eXpression (NPX) values, representing relative protein abundance on a log₂ scale, were generated using Olink’s quality control workflow. Full details on protein measurement and QC are provided in ref^20^. Nightingale Health NMR metabolomics data (Category 220) were derived for 251 metabolites (including ratios and percentages) (average N=487K), which includes lipoproteins, fatty acids, free cholesterol and amino acids. Metabolite concentrations were IRNT normalized prior to analysis. Dietary phenotypes were derived from the UKB touchscreen questionnaire (Category 100052). We selected 20 dietary variables spanning fat-related intake (e.g., pork: Field 1389, beef: Field 1369, lamb/mutton: Field 1379), carbohydrate-related intake (e.g., bread: Field 1438), and additional control phenotypes (**Supplementary Table 4**). For all phenotypes, analyses were restricted to the baseline assessment (i0: 2006-2010) to ensure temporal alignment across data modalities. All UK Biobank analyses were performed on the Research Analysis Platform (RAP) under application 16549.

### stressor Protein Program (sPP) scores

To quantify the activity of stressor gene programs in circulating plasma, we computed stressor Protein Program (sPP) scores using protein abundance data from the UK Biobank Pharma Proteomics Project (UKB-PPP). The UKB-PPP measured plasma concentrations of 2,819 proteins using the Olink Explore platform across 45,956 White British individuals. For each stressor program, we mapped DEGs to the Olink protein panel by matching gene symbols to their corresponding protein products. Let *G* denote a stressor gene program and *G*′ ⊂ *G* denote the subset of program genes with corresponding proteins measured in the Olink panel. We used AUCell^48^ to compute per-individual enrichment scores for each plasma-mapped stressor program. AUCell quantifies gene set activity based on the recovery of program genes within the top-ranked features of each sample, providing a robust measure of coordinated program expression that is not sensitive to absolute expression scale. First, all 2,819 proteins are ranked by their normalized abundance (NPX) in individual *i*, from highest to lowest. Let *r_ip_* denote the rank of protein *p* in individual *i*. For the protein-mapped gene set *G*’, we compute a cumulative recovery curve; at each position *x* in the ranking (from 1 to the total number of proteins), we calculate the fraction of genes in *G*’ recovered

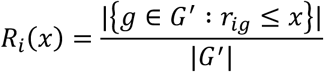

The AUCell score is a continuous score obtained by taking the area under this curve. The resulting AUC value for individual *i* and stressor program *G*^0^ is denoted as the stressor Plasma Proteome score, *sPP*_*i*_(*G*’). The AUC ranges from 0 to 1, where higher values indicate that program genes are concentrated among the most highly expressed proteins in that individual. We computed sPP scores for each stressor condition separately for uDEG and dDEG programs.

Because plasma integrates secretory output across tissues, sPP scores reflect β-cell stress program activity as read out through program-relevant proteins that reach circulation; not all program genes are represented in plasma, and program proteins that are well-represented may also be contributed by non-pancreatic tissues. We interpret sPP scores as systemic readouts of β-cell stress program state, with genetic regulation potentially acting through either direct β-cell mechanisms or trans-tissue pathways that shape the circulating environment.

### Diet-tertile stratification analysis of the sPP scores

To characterize associations between dietary intake patterns and sPP scores, we performed tertile-based stratification analyses across dietary phenotypes. For each dietary phenotype, individuals were stratified into tertiles (low, medium, high) based on intake frequency or quantity. Ties at tertile boundaries were assigned to the middle tertile. We computed the mean sPP score (post quantile matching across sPPs) within each tertile and visualized the relationship between dietary exposure and sPP activity using 1D heatmaps. To quantify the magnitude of sPP variation against dietary exposure *D*, we defined the 1D dietary gradient score as follows:

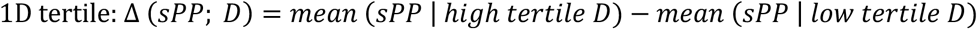

Besides these 1D tertile maps, to assess joint effects of dietary exposures, we performed 2D tertile mapping for biologically relevant diet pairs (e.g., bread intake × pork intake, representing carbohydrate and fat intake axes). Individuals were cross-classified into 9 cells based on tertile membership for each of the two dietary variables (*D*_1_, *D*_2_ ∈ {*Low*, *Medium*, *High*}), and the mean sPP score was computed within each cell. For the 2D tertile map of dietary combinations, we define high-diet-combo cells (*C*^+^) as those satisfying max(*D*_1_, *D*_2_) = High and min(*D*_1_, *D*_2_) ≥ Medium (High/High, High/Medium, Medium/High), and low-diet-combo cells (*C*^3^) satisfying min(*D*_1_, *D*_2_) = Low. The 2D dietary gradient score was computed as:

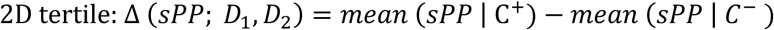

Positive values of Δ for both 1D and 2D tertile maps demonstrates that the sPP score is positively associated with higher intake and joint intake of dietary variables. Significance of 1D and 2D tertile gradient scores was assessed by standard 2-sample t-test between the high and low tertile groups as reported above.

### Genome-wide sPP-QTL mapping

We performed genome-wide testing of 9,124,756 imputed autosomal variants (MAF ≥ 0.01) against 40 sPP scores based on plasma protein and genotyping data from 45,956 White British individuals from UK Biobank. sPP scores were rank normalized to a standard normal distribution prior to association testing. Association testing between sPP scores and allelic dosage was performed using REGENIE (v4.1.2), a two-step linear mixed model framework that accounts for population structure and relatedness^94^. We used as covariates age, sex, age^2^, age × sex, the first 10 genetic principal components, and genotyping array. In Step 1, REGENIE employs a whole-genome ridge regression model on a filtered set of directly genotyped variants to generate leave-one-chromosome-out (LOCO) polygenic predictions for each individual. Variants are filtered based on MAF > 1%, genotyping rate per variant >99%, and genotyping rate per individual >80%. In Step 2, we tested imputed and QC-ed variants (per-variant genotyping rate < 99% or imputation INFO score < 0.7) for association with sPP scores using linear regression, with the LOCO predictions used to account for polygenic background effects. Genome-wide significant loci (P < 5e-08) were defined by merging overlapping 1 Mb windows around each significant variant into distinct genomic regions. Each locus was annotated by the lead variant with the strongest association. Lead variants were annotated to the nearest gene by transcription start site (TSS) distance and classified as *cis*-acting if located within 1 Mb of a gene encoding a protein in the corresponding sPP score, and *trans*-acting otherwise. Genomic inflation was assessed using the genomic control factor (*λ*_*GC*_) from REGENIE and the LD score regression (LDSC) intercept^90^. SNP-based heritability (*h_g_*^2^) was estimated by applying LDSC to GWAS summary statistics using the 1000 Genomes European LD reference panel. Regional association plots were generated using LocusZoom^95^.

### Fine-mapping and colocalization of sPP-QTL

To identify putative causal variants at genome-wide significant sPP-QTL loci, we applied statistical fine-mapping using the SuSiE (Sum of Single Effects) model^96^. For each independent locus, we defined a fine-mapping region as a 1 Mb window centered on the lead variant. In-sample linkage disequilibrium (LD) matrices were previously computed from UK Biobank White British individuals^97^. SuSiE was run with a maximum of 10 causal signals per locus (L = 10) and default prior variance settings. For each independent signal, we report 95% credible sets and variant-level posterior inclusion probability (PIP). A variant with PIP>0.2 are denoted as “likely causal” variants for the associated sPP scores.

To assess whether sPP-QTL signals share causal variants with T2D and related cardiometabolic traits, we performed multi-phenotype colocalization analysis using ColocBoost^59^. For each of 104 UK Biobank complex diseases and traits (**Supplementary Table 5**), we performed disease-targeted ColocBoost with the disease/trait summary statistics as the target and 40 sPP-QTL summary statistics as secondary phenotypes. To avoid any bias, we defined analysis windows as ±1 Mb around gene transcription start sites rather than centering on disease GWAS or sPP lead variants. ColocBoost was applied using LD reference panels computed from UK Biobank White British individuals^97^. We report two metrics from ColocBoost: (i) colocalization events, each comprising a 95% colocalization set (CoS) of variants tied to a set of colocalizing phenotypes, and (ii) variant-level VCP (variant colocalization probability) scores quantifying confidence in shared causal variants within each CoS. **Supplementary Figure 6f** presents a summary of 104 ColocBoost runs, each containing a disease/trait GWAS from UK Biobank against 40 sPP traits; we plot the number of CoS corresponding to each stressor (row) and disease/trait (column) (**Supplementary Table S5).**

### sPP variance QTL (sPP-vQTL) mapping

To identify genetic variants associated with inter-individual variance in sPP scores, we performed variance QTL (vQTL) mapping using OSCA^64^, following the strategies adopted in refs^65,66,98^. Variance effects can indicate context-dependent genetic architecture, where a variant’s effect on the phenotype depends on unmeasured environmental, physiological, or metabolic contexts—a hallmark of genotype-by-environment (G x E) interaction. We applied the Bartlett and Levene test, which is robust to departures from normality. In Levene’s test, individuals were stratified by genotype (0, 1, or 2 copies of the effect allele), and the absolute deviation from the genotype-group median was computed for each individual. Association between genotype and absolute deviation was tested using linear regression, adjusting for the same covariates as the mean-effect GWAS (age, sex, age^2^, age × sex, 10 genetic PCs, and genotyping array). Both tests were applied to the glucolipotoxicity sPP score across 45,956 UK Biobank White British individuals. To distinguish true variance effects from artifacts driven by mean-variance relationships, we excluded variants with genome-wide significant mean effects (*P_mean_* < 5e-08) from vQTL interpretation, as scale-dependent mean-variance coupling can produce spurious variance association. Given that non-spurious vQTL effects are expected to be weaker compared to mean effect changes, we used a more flexible genome-wide significance cut-off of 1e-05 to call a vQTL association as significant. To further characterize the metabolic contexts driving variance effects, lead vQTL variants were tested for explicit genotype-by-environment interactions.

### G x Diet interaction test

Self-reported intake from the UK Biobank touchscreen food-frequency questionnaire (FFQ) is subject to substantial measurement error, recall bias, and a heavy right tail, and the set of variants carried forward for gene–diet (G x diet) testing was deliberately small. We therefore modelled dietary intake as a *discretised* exposure and assessed interaction with a single omnibus contrast test per locus rather than a continuous interaction coefficient. For each focal diet, participants were partitioned into low-, medium-, and high-intake tertiles of the intake distribution. We propose an omnibus contrast test (OC) that retains power against both monotone and non-monotone effect-modification patterns without pre-committing to a directional hypothesis.

Let *y*_*i*_ denote the quantile-matched nutrient-stress sPP scores (facilitating robust comparisons across stress programs) for individual *i*, *g*_*i*_ is the allele dosage (0, 1, 2) at the focal typed variant, and *k*(*i*)*∈*{1,2,3} the assigned intake tertile. We code the ordered three-level exposure with orthogonal polynomial contrasts — a linear basis *ϕ*^(1)^ ∝ (−1,0, +1) and a quadratic basis *ϕ*^(2)^ ∝ (+1, −2, +1) — assigning each individual the corresponding polynomial scores *P_i_*^(1)^and *P_i_*^(2)^. The interaction model is

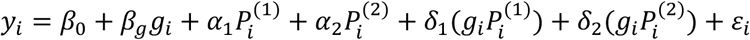

where (*δ*_1_, *δ*_2_) capture the linear and quadratic components of how the per-allele effect of *g*_*i*_ on the sPP score varies across the dietary gradient. The **omnibus contrast test** is the joint null *H*_0_: *δ*_1_ = *δ*_2_ = 0, evaluated as

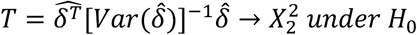

with *var*(*δ̂*) a heteroskedasticity-consistent (HC3 sandwich) estimator to guard against residual within-tertile variance heterogeneity that survives the global RINT. This approach is algebraically equivalent to the 2-df joint test of a genotype × tertile-indicator interaction. To prevent spurious interaction signal arising from scale and confounding artefacts, the sPP score was RINT transformed, and adjusted for age, sex, age^2^, age × sex, the first 10 genetic principal components, and genotyping array.

### Cross Diet Genotype Alignment (CDGA)

To quantify the concordance of genotype dosage effects across the lipid and glucose dietary arms of glucolipotoxicity, we defined a Cross Diet Genotype Alignment (CDGA) index for each variant. For each SNP *s*, we extracted average quantile-matched sPP scores of individuals corresponding to four cells of the genotype × diet interaction grid: the homozygous reference (0/0) and homozygous alternate (1/1) genotype groups, each stratified by the lowest (Low) and highest (High) tertile of macronutrient intake, separately for pork intake (representing lipid load) and bread intake (representing carbohydrate load). Variants were retained for analysis only if each of the four corner cells contained at least 1,000 individuals in both the pork and bread strata, to ensure we have sufficient power to detect genotype specific effects given these effects are expected to be weak.

For each retained variant, we computed the genotype effect of SNP *s* on sPP scores at each diet extreme *d* ∈ {*Low*, *High*} for macronutrient arm *m* ∈ {*pork*, *bread*}.

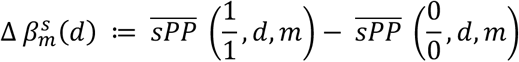

The CDGA index for SNP *s* is then defined as the sum of cross-diet products of these dosage-effect terms across the two dietary extremes.

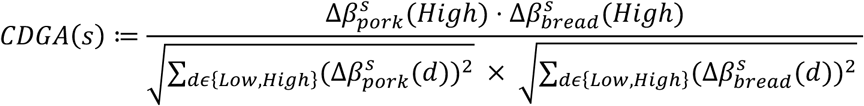

### Effect Direction Concordance and enrichment metrics

To quantify the extent of overlap between two distinct DE gene sets relative to the full set of genes, we calculate the odds ratio (OR). To ensure numerical stability, we add a pseudocount of 5 to each cell of the contingency table in all cases prior to computing the OR. To evaluate whether the observed enrichment is statistically significant (i.e., whether the OR is significantly greater than 1), we performed Fisher’s exact test.

Furthermore, to assess the concordance in the direction of expression changes between two DE gene sets, we define the effect direction concordance (EDC) as follows:

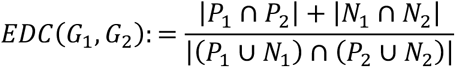

where *P*_1_ (*N*_1_) and *P*_2_ (*N*_2_) denote the sets of genes with positive (negative) log fold changes in DE gene sets *G*_1_and *G*_2_, respectively. This metric represents the fraction of shared DE genes that exhibit concordant directions of change. We test whether this fraction exceeds 0.5 using a binomial test.

## Supplementary Figures

**Supplementary Figure 1:**
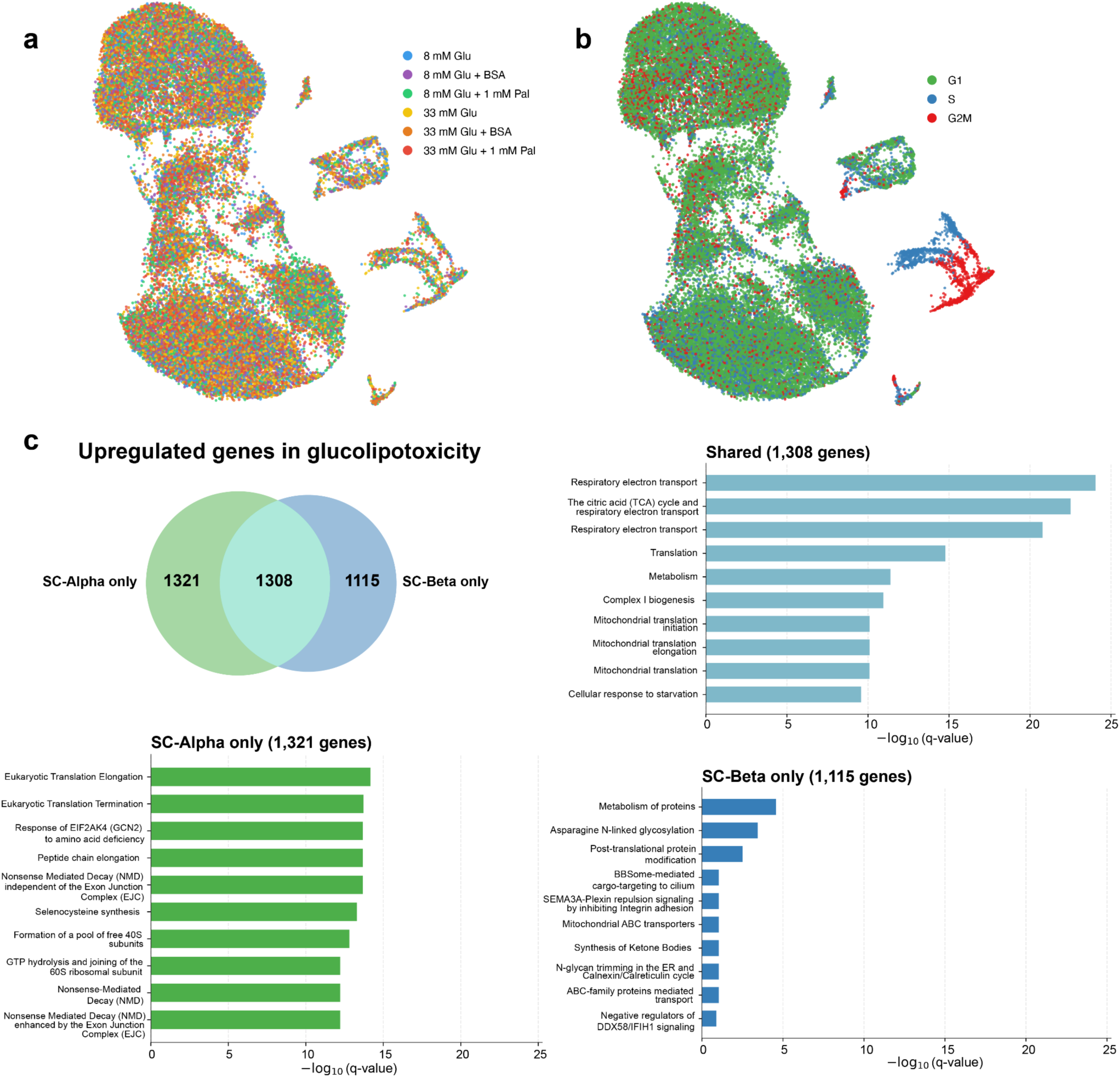
Glucolipotoxicity induces shared and cell type–specific transcriptional programs in SC-α and SC-β cells. **a.** UMAP of all profiled SC-islet cells (N = 38,350) colored by nutrient-stress condition (8 mM glucose; 8 mM glucose + BSA; 8 mM glucose + 1 mM palmitate; 33 mM glucose; 33 mM glucose + BSA; 33 mM glucose + 1 mM palmitate). **b.** The same UMAP as in (a) colored by inferred cell-cycle phase (G1, S, and G2/M). **c.** Shared and cell-type-specific glucolipotoxicity-upregulated genes (uDEGs) in SC-á and SC-β cells. Venn diagram showing the extent of this sharing. Bar plot representing top pathways from pathway enrichment analysis performed on (i) shared uDEGs of glucolipotoxicity in SC-α and SC-β cells, (ii) glucolipotoxicity uDEGs specific to SC-α cells, and (iii) glucolipotoxicity uDEGs specific to SC-β cells. Numerical results are reported in **Supplementary Table S1**.

**Supplementary Figure 2:**
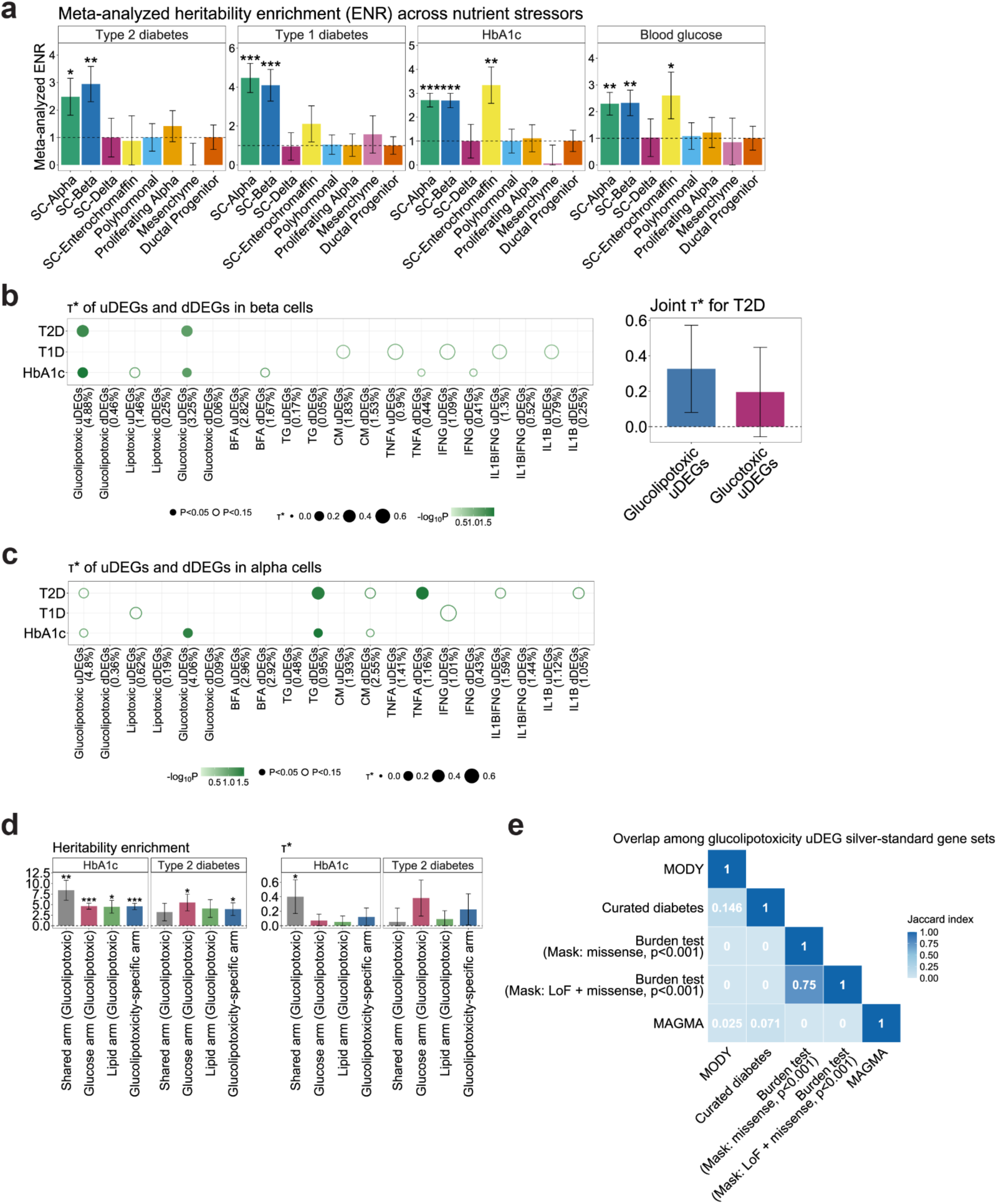
Additional results from comparative analysis of stress-induced signatures. **a.** Meta-analyzed heritability enrichment (ENR) across all nutrient stress-induced uDEGs and dDEGs for each related trait (T2D, T1D, HbA1c, and blood glucose) and cell type (SC-Alpha, SC-Beta, SC-Delta, SC-Enterochromaffin, Polyhormonal, Proliferating Alpha, Mesenchyme, and Ductal Progenitor). In the meta-analysis, heritability enrichment and SE were set to 1 for cases with fewer than 100 uDEGs or dDEGs due to unreliable estimates (actual estimates in **Supplementary Table S2**). **b.** Left: Standardized effect sizes (*τ*\*) of uDEGs and dDEGs in β cells across 10 stressors (nutrient + ER + inflammatory) for T2D, T1D, and HbA1c. Dot size represents the magnitude of *τ*\*, dot color represents significance (−log₁₀P), filled dots denote P < 0.05 and open dots P < 0.15. The percentage in parentheses below each DEG set name indicates the proportion of common variants falling within the regulatory annotation (linked ENCODE-rE2G-max enhancers) of that gene set. Right: T2D *τ*\* in a joint model comprising glucolipotoxicity uDEGs and glucotoxicity uDEGs, both of which are marginally significant; glucolipotoxicity absorbs the heritability signal from glucotoxicity in the joint model. **c.** Standardized effect sizes (*τ*\*) of uDEGs and dDEGs in α cells across the same 10 stressors for T2D, T1D, and HbA1c. Dot size, color, and shape follow the same conventions as in panel **b**. This plot highlights that T2D heritability informativeness of glucolipotoxicity and glucotoxicity uDEGs is specific to SC-β cells. **d.** T2D and HbA1c heritability enrichment (ENR) and standardized effect sizes (*τ*\*) of 413 uDEGs shared across all three nutrient stressors (shared arm), 688 uDEGs shared exclusively between glucolipotoxicity and glucotoxicity uDEGs (glucose arm), 151 uDEGs shared exclusively between glucolipotoxicity and lipotoxicity uDEGs (lipid arm), and 1,171 glucolipotoxicity-specific uDEGs (glucolipotoxicity-specific arm) in SC-β cells. **e.** Pairwise overlap (Jaccard index) of five diabetes-related gene sets intersecting with glucolipotoxicity uDEGs in SC-β cells. The five gene sets include MODY genes, curated diabetes genes, and genes identified through rare-variant burden tests using two coding variant masks (missense and LoF + missense; P < 0.001), and MAGMA-prioritized genes (Z-score > 3) for T2D. Asterisks in panels **a** and **d** denote significance from a one-sided Z-test assessing whether meta-analyzed ENR > 1 and ENR > 1, respectively (*** P < 0.001, ** P < 0.01, * P < 0.05). Error bars in **a**, **b**, and **d** represent ±1SE. Numerical results are reported in **Supplementary Table S2**.

**Supplementary Figure 3:**
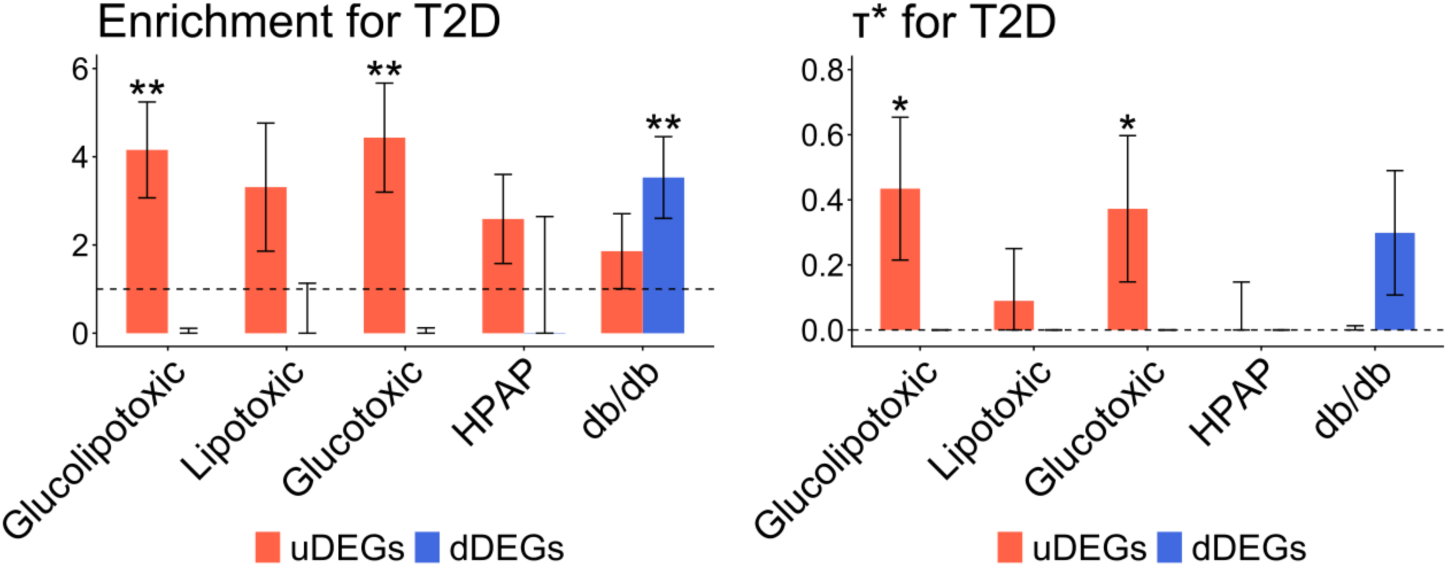
T2D heritability enrichment after calibrating DEG counts between nutrient stress and other in-vivo islet DEGs. T2D heritability enrichment (ENR; left) and standardized effect sizes (*τ*\*; right) for uDEGs (red) and dDEGs (blue) from three nutrient-stress programs (SC-β cells), HPAP, and db/db (primary β cells). To match the DEG count of glucolipotoxicity uDEGs (n = 2,423), HPAP and db/db DEGs were subsampled to the top 2,500 DEGs ranked by P-value when the number of DEGs exceeds 2,500. Error bars represent ±1 SE. Asterisks denote significance from a one-sided Z-test assessing whether ENR > 1 or *τ*\* > 0 (*** P < 0.001, ** P < 0.01, * P < 0.05). Numerical results are reported in **Supplementary Table S3**.

**Supplementary Figure 4:**
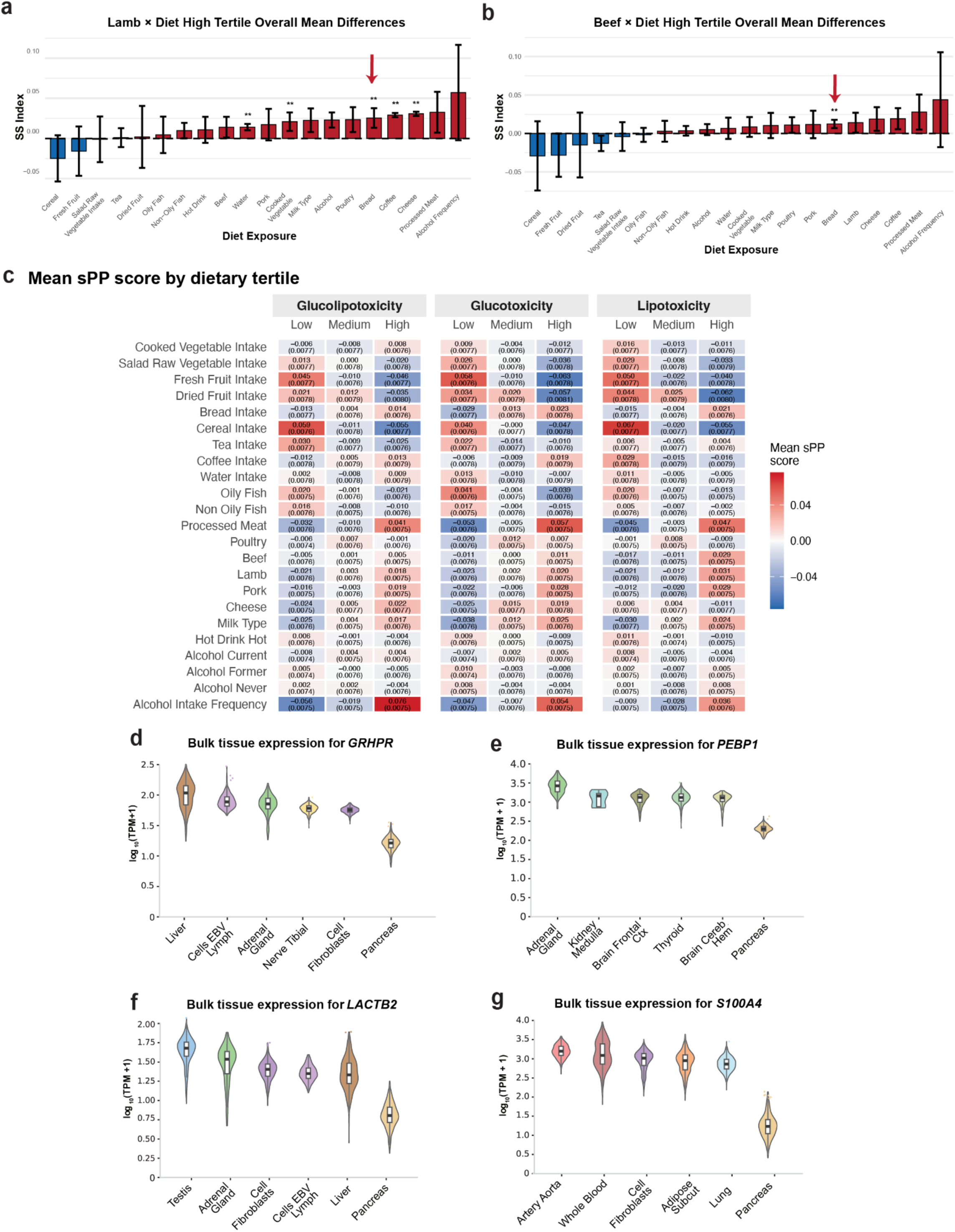
Additional dietary association map of glucolipotoxicity uDEG sPP score. **a,b**. Bidirectional dietary-specificity analysis for glucolipotoxicity sPP with red-meat intake held high. For lamb (**a**) and beef (**b**), each bar shows, for the indicated second dietary exposure, the difference between the mean glucolipotoxicity sPP across the high-intake tertile combinations [(High, Medium), (High, High), (Medium, High)] and the overall mean. Bread (red arrow) shows the strongest positive difference among all second exposures, supporting the carbohydrate × saturated-fat combination as the driver of the signature. Error bars represent ±1 SE; asterisks denote significance (*** P < 0.001, ** P < 0.01, * P < 0.05). **c.** Mean glucolipotoxicity, glucotoxicity, and lipotoxicity sPP scores across Low/Medium/High tertiles of each dietary exposure. Each cell shows the mean sPP score (SE in parentheses), colored from low (blue) to high (red). **d-g**. GTEx bulk-tissue expression (log₁₀[TPM + 1]) of the top non-program plasma proteins correlated with glucolipotoxicity sPP: GRHPR (**d**), PEBP1 (**e**), LACTB2 (**f**), and S100A4 (**g**) — showing the 5 highest-expressing tissues, with pancreas included for reference. Each protein is expressed predominantly in extra-pancreatic tissues (liver, adrenal gland, adipose, and others) rather than pancreas, consistent with their origin as non-program, cross-tissue correlates of β-cell stress activity in plasma. Numerical results are provided in **Supplementary Table S4**.

**Supplementary Figure 5:**
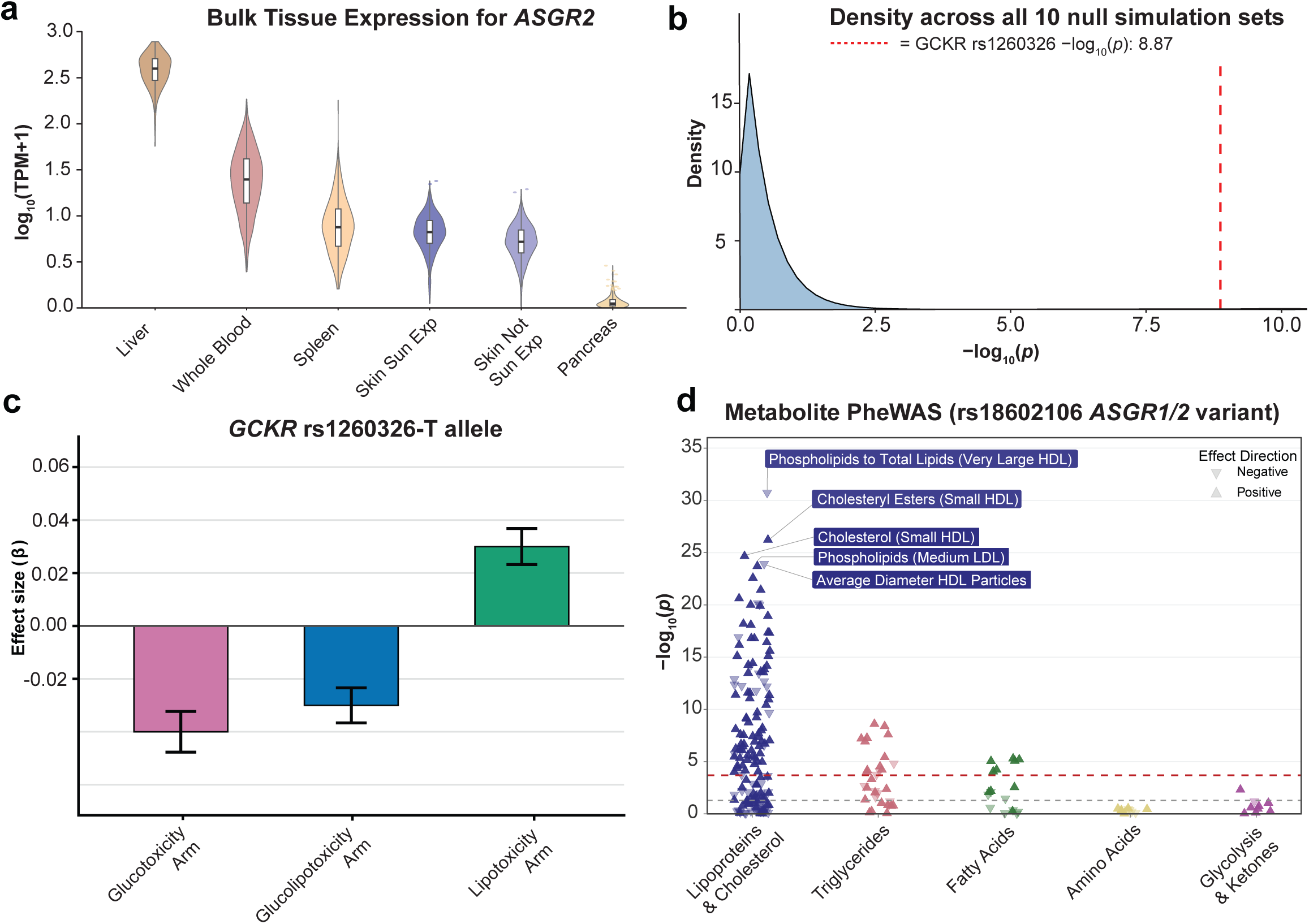
Additional vignettes around genetic variants at *GCKR* and *ASGR1/2* loci associated with glucolipotoxicity uDEG sPP. **a.** GTEx bulk-tissue expression (log₁₀[TPM + 1]) violin plot of *ASGR2* across its 5 highest-expressing tissues (pancreas shown for reference). Expression is predominantly liver-restricted, supporting a hepatic origin for the *ASGR1/2* trans-sPP-QTL signal. **b.** Specificity of the *GCKR* rs1260326 association with glucolipotoxicity sPP. Density of −log₁₀(P) values for *rs1260326* against 100 size-matched null sPP scores across null simulations of randomly sampled sets of proteins of the same size as the glucolipotoxicity uDEGs. The original −log₁₀(P) for glucolipotoxicity uDEG sPP at this variant is highlighted by a vertical red dotted line. **c.** Effect of the *rs1260326* T allele (β ± SE) on the three arm-specific glucolipotoxicity sPP scores — glucose arm, glucolipotoxicity-specific arm, and lipid arm. **d.** Metabolite PheWAS (−log₁₀ P) of the *ASGR1/2* variant *rs186021206* across NMR metabolite classes. Positive and negative associations are denoted by upward and downward triangles. The dashed line denotes the Bonferroni significance cut-off. The metabolites are grouped into five classes (lipoproteins & cholesterol, triglycerides, fatty acids, amino acids, glycolysis & ketones). Numerical results are provided in **Supplementary Table S5**.

**Supplementary Figure 6:**
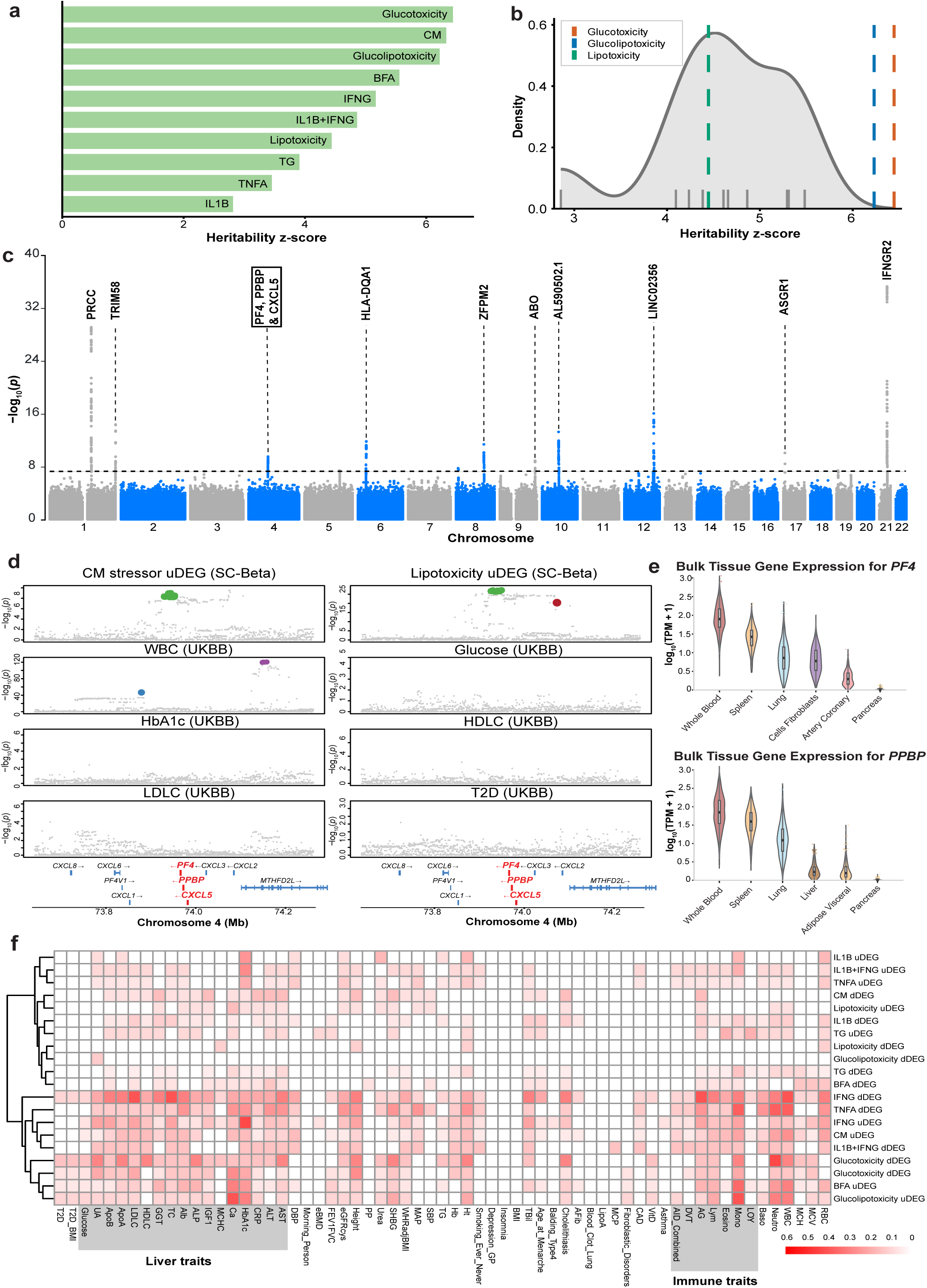
Genome-wide heritability, null calibration, and trans-tissue / multi-phenotype regulation of stress-program sPP-QTLs. **a.** SNP-based heritability z-scores (LDSC) for the stressor sPP uDEG scores (nutrient, ER, and inflammatory programs) in β cells. Glucotoxicity, cytokine-mix (CM), glucolipotoxicity, and BFA show the highest heritability. **b.** Heritability z-scores for null sPP scores generated from random size-matched protein sets (gray density), compared with the three nutrient-stress sPPs (glucotoxicity, glucolipotoxicity, lipotoxicity; colored dashed vertical lines). **c.** sPP-QTL Manhattan plot of the cytokine mix (CM) uDEGs in primary β cells. **d.** Regional multi-phenotype colocalization (using ColocBoost) at the *PF4/PPBP/CXCL5* chemokine cluster locus between cytokine mix (CM) uDEG sPP-QTLs in primary β cells, lipotoxicity uDEG sPP-QTLs in SC-β cells, and other GWAS traits of interest. **e.** GTEx bulk-tissue expression (log₁₀[TPM + 1]) of *PF4* and *PPBP*, highest in whole blood and spleen and minimal expression in pancreas, consistent with an immune-tissue origin for the chemokine-cluster sPP-QTL. **f.** Multi-phenotype colocalization between stressor sPP-QTLs (rows; β cells up- and down-regulated DEG sets across stressors, hierarchically clustered) and 104 UK Biobank complex traits (columns, grouped — including liver and immune traits). Cell color indicates the fraction of colocalizations observed with our sPP-QTLs and trait of interest among all putative causal variants for the sPP-QTLs. Numerical results are provided in **Supplementary Table S5.**

**Supplementary Figure 7:**
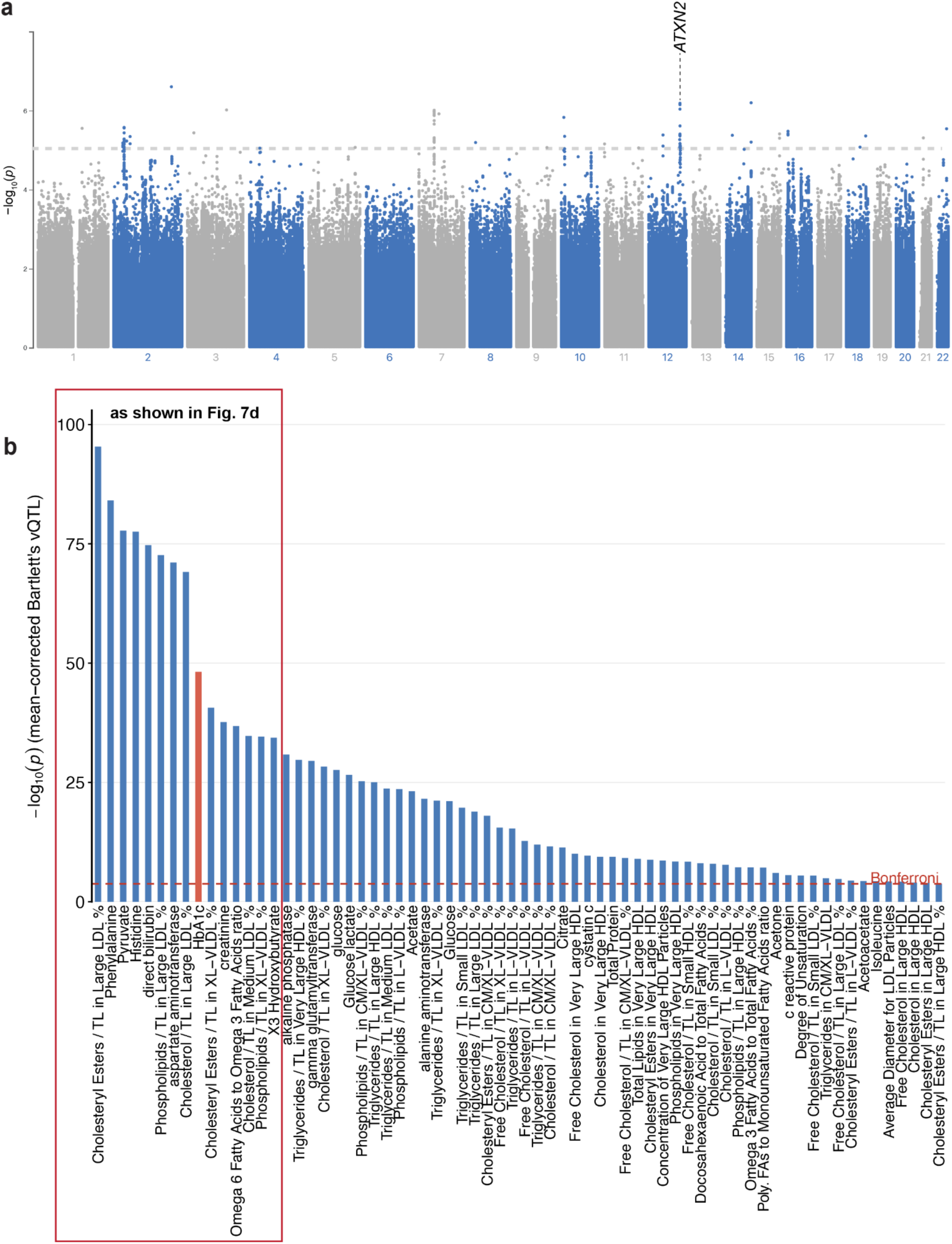
Variance-QTL (vQTL) analysis at *ATXN2/SH2B3* locus (*rs10774625*). **a.** Bartlett’s vQTL association (−log₁₀ P) on RINT-transformed glucolipotoxicity uDEG sPP identifying rs10774625 in the *ATXN2/SH2B3* locus as the sentinel variance-QTL hit. The dashed line denotes the suggestive 1e-05 cut-off. **b.** vQTL metabolite PheWAS: Bartlett’s variance-QTL associations (−log₁₀ P) of RINT-transformed NMR circulating metabolites and blood biomarker traits at the *rs10774625* variant, ordered by significance. The horizontal red dotted line corresponds to Bonferroni significance cutoff. Numerical results are reported in **Supplementary Table S6.**

